# Heritability within groups is uninformative about differences among groups: cases from behavioral, evolutionary, and statistical genetics

**DOI:** 10.1101/2023.11.06.565864

**Authors:** Joshua G. Schraiber, Michael D. Edge

**Affiliations:** Department of Quantitative and Computational Biology, University of Southern California

## Abstract

Without the ability to control or randomize environments (or genotypes), it is difficult to determine the degree to which observed phenotypic differences between two groups of individuals are due to genetic vs. environmental differences. However, some have suggested that these concerns may be limited to pathological cases, and methods have appeared that seem to give—directly or indirectly—some support to claims that aggregate heritable variation within groups can be related to heritable variation among groups. We consider three families of approaches: the “between-group heritability” sometimes invoked in behavior genetics, the statistic *P*_*ST*_ used in empirical work in evolutionary quantitative genetics, and methods based on variation in ancestry in an admixed population, used in anthropological and statistical genetics. We take up these examples to show mathematically that information on within-group genetic and phenotypic information in the aggregate cannot separate among-group differences into genetic and environmental components, and we provide simulation results that support our claims. We discuss these results in terms of the long-running debate on this topic.

## 1 Introduction

Understanding the causes and consequences of phenotypic variation is a central goal of many fields of research. Over the last century, researchers in multiple fields have attempted to make a study of the genetic and environmental variation within groups of organisms and use that information to understand the genetic and environmental sources of among-group differences. Such attempts have included suggestions that the explanation of phenotypic differences within groups likely extends to differences among groups (Jensen, 1969), suggestions that observed differences may result from adaptation (for example, Castellani et al., 2023; Salloum et al., 2022; Vitek et al., 2022), and observations that admixed individuals have phenotypes intermediate with respect to members of source populations (Chan et al., 2015; Fernández et al., 2003; Kosoy et al., 2012; Molokhia et al., 2003).

Richard Lewontin critiqued the first of these arguments in a set of well-known thought experiments (1970). In one example, Lewontin imagined two groups of corn plants. Within each group, genotypes vary, but environmental conditions do not. Between groups, there are no average genetic differences, but environmental conditions differ, with one group in more favorable conditions than the other. In this situation, within-group phenotypic variation is genetic (since environmental conditions are the same for all members of the same group), while between-group variation is environmental (since there are no average genetic differences between groups). Lewontin claimed that his thought experiments showed that within-group heritability is irrelevant for determining the causes of among-group differences. Lewontin’s argument was a response to Arthur Jensen, who claimed that within-group heritability estimates suggested a genetic explanation for racial differences in scholastic achievement (Jensen, 1969).

In the decades since Lewontin wrote, several researchers have suggested that Lewontin’s thought experiment represents a pathological special case. For example, in behavioral genetics, DeFries (1972), replying to both Lewontin and Jensen, built on earlier work by Lush (1949) to propose the “between-group heritability,”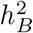, aiming to quantify the relative influence of genetic and environmental factors on phenotypic differences among groups. DeFries argued that Lewontin’s thought experiments correspond to specially picked edge cases and suggested that in other cases,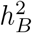 provides a useful framework for thinking about group difference.

Here, we show that in contrast to DeFries’ suggestion, the rationale underlying Lewontin’s thought experiments is general and poses challenges for the interpretation of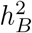whenever environmental differences among groups are not controlled or understood. In particular, we show that, in line with Lewontin’s intuition, the heritabilities of phenotypic differences between groups are not constrained by heritability within groups: perfect knowledge of within-group heritability provides no information about between-group heritability. Crucially, even if the heritability of between-group differences is estimated correctly, it leaves the direction of the genetic and environmental components of phenotypic difference unclear.

We also expand Lewontin’s critique to other approaches to similar questions. In evolutionary biology, a statistic called *P*_*ST*_ (Brommer, 2011) is used to test whether phenotypic differences among groups of wild organisms are consistent with neutral evolution or instead might be better explained by natural selection. We show that a parameter used in sensitivity analyses involving *P*_*ST*_ can be understood as a function of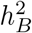. Using our results about 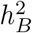and their relationship to *P*_*ST*_, we show the sensitivity analysis that is typically under-taken when using *P*_*ST*_ to test for evidence of divergent selection on a phenotype may be conservative, especially if the phenotypic differentiation among groups is small. Moreover, we highlight the difficulty of interpreting *Q*_*ST*_ and *P*_*ST*_ in the presence of environmental differences between groups, adding to recent arguments by Harpak and Przeworski (2021).

In genetic anthropology and statistical genetics, variation in ancestry among admixed individuals is sometimes used to make claims about the sources of trait variation among groups. For example, correlations between an individuals’ phenotype and ancestry proportions may be taken as evidence that an observed phenotypic difference between source populations is genetically caused. As we show and has been mentioned previously as a cautionary point (e.g., Chakraborty & Weiss, 1986; Fernández et al., 2003; Kosoy et al., 2012), the genetic effect of ancestry may be confounded with environmental variation affecting the phenotype. Approaches to heritability estimation based on patterns of local ancestry in admixed populations (Zaitlen et al., 2014)—as opposed to global ancestry proportions— have also been used to calibrate claims about the sources of group differences (Zaidi et al., 2017). We show that the variation due to local ancestry that they leverage has no clear relationship with between-group trait differences.

We discuss our results in terms of their shared origin in gene-environment confounding and relate them to previous claims (Block, 1995; Feldman & Lewontin, 1975; R. Lewontin, 1970). The complications we point out arise under simple additive models, and the situation is only more complicated in models with interactions among genetic or environmental influences. We defer a brief consideration of gene-by-environment interaction to the discussion section.

## 2 Between-group heritability is unconstrained by within-group heritability and underdetermines genetic and environmental differences among groups

Lush (1949) developed the between-group heritability,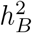, to quantify the relative influence of genetic and environmental factors on a trait that differs among groups. Lush’s interest was in heritability of differences among families of organisms subject to artificial selection, with the goal of weighting family membership appropriately when selecting organisms for breeding. Lush worked in applications in which many aspects of among-group variation in environment could be controlled or randomized.

Consider a phenotype value *Y* formed from additive genetic and environmental components, i.e. *Y*_*i*_ = *G*_*i*_ + *E*_*i*_ for individual *i*. It is standard to define the (narrow-sense) heritability as *h*^2^ = *V*_*A*_*/V*_*P*_, where the additive genetic variance *V*_*A*_ is the variance of the genetic component (*G*_*i*_), assuming no epistasis or dominance, and *V*_*P*_ is the variance of the phenotype. If the sample is divided into groups, we can also define variance proportions accounted for by group membership in either traits themselves or in the genetic components of those traits. For example, with respect to the phenotypic variance, we can define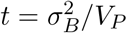, with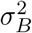 the between-group component of the phenotypic variance (Falconer & Mackay, 1996). Call the analogous variance proportion for the genetic variance *r*. Then, when groups are large, the “between-group” heritability is

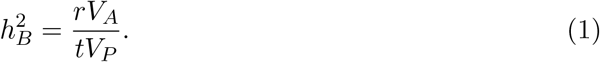

Subsequently, DeFries (1972) took up this formalism to argue against the generality of Lewontin’s thought experiment. With some algebra, he showed that the between-group heritability is mathematically related to the within-group heritability,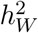. Specifically, defining *r* and *t* as in the previous paragraph, he showed that

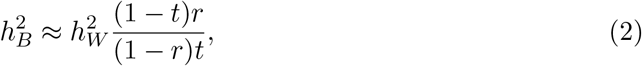

if the groups are large. DeFries claimed that Lewontin’s thought experiments—one described above, and another involving inbred lines reared in different environments—could be explained as pathological cases in which either *r* or *t* is equal to 0, and suggested that the relationship in equation (2) could be used to help interpret observed group differences in other cases. He displayed a table showing between-group heritabilities implied by various values of *r*, conducting a sort of sensitivity analysis.

An immediate problem with application of equation (2) is that in the absence of environmental controls, it is impossible to estimate *r*, the proportion of the variance in the genetic component of the trait accounted for by group membership, a point that DeFries conveyed (DeFries, 1972). Despite this, some researchers in behavior genetics, including DeFries himself, took up DeFries’ formalism and argued for its relevance for understanding variation among groups of humans, generally in the context of discussions of racial differences in IQ or scholastic achievement (Plomin & DeFries, 1976; Warne, 2020, 2021). In particular, Jensen (1972a, 1972b, 1975) repeatedly cited the definition of between-group heritability as a linear function of within-group heritability in arguments for the value of heritability estimates for understanding sources of difference among groups. Feldman and Lewontin (1975) criticized the expression for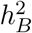 in equation 2 as tautological, in that *r* is better viewed as a term that simply relates the definitions of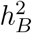and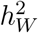 rather than being an independently varying ingredient of a prediction of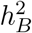 from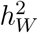. Here, we concur with Feldman and Lewontin’s interpretation, showing that, despite the appearance of equation (2), the between-group heritability is unconstrained by the within-group heritability. Thus, within-group heritability, on its own, provides no information about between-group heritability (Supplementary Text, section S1.1-S1.2). We also show that even if *r* is known, 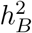 still underdetermines the group differences, such that opposing interpretations of among-group variation are possible under the same (known) *r* and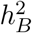.

In the specific case of two groups, the important quantities are *δ*_*G*_, the difference between the mean genetic values of the two groups, and *δ*_*E*_, the difference between the mean environmental values of the two groups. Then, under a standard quantitative-genetic model,

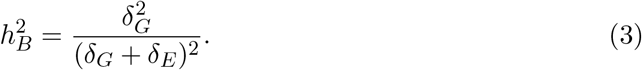

Equation (3) is useful to highlight several properties of between-group heritability that generalize to cases with more than two groups. First, this expression for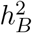 can be derived without reference to within-group heritability or any quantities that contribute to within-group heritability (section S1.2), indicating that even perfect knowledge of heritability within groups conveys no information about genetic differences between groups without further information or assumptions.

Second, it shows that the between-group heritability is not bounded between 0 and 1: in particular, if the environmental and genetic effects point in different directions, such that the magnitude of the phenotypic deviation, *δ* = *δ*_*G*_ + *δ*_*E*_ is smaller than the magnitude of the genetic deviation, *δ*_*G*_, then the between-group heritability will be larger than 1. We illustrate this in Figure 1A, which shows the value of 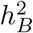across a range of genetic and environmental deviations. One striking feature of the behavior of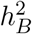 is that it rapidly approaches infinity when *δ*_*E*_ *≈ −δ*_*G*_, and that larger genetic deviations result in a larger range of environmental deviations over which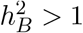.

**Figure 1.**
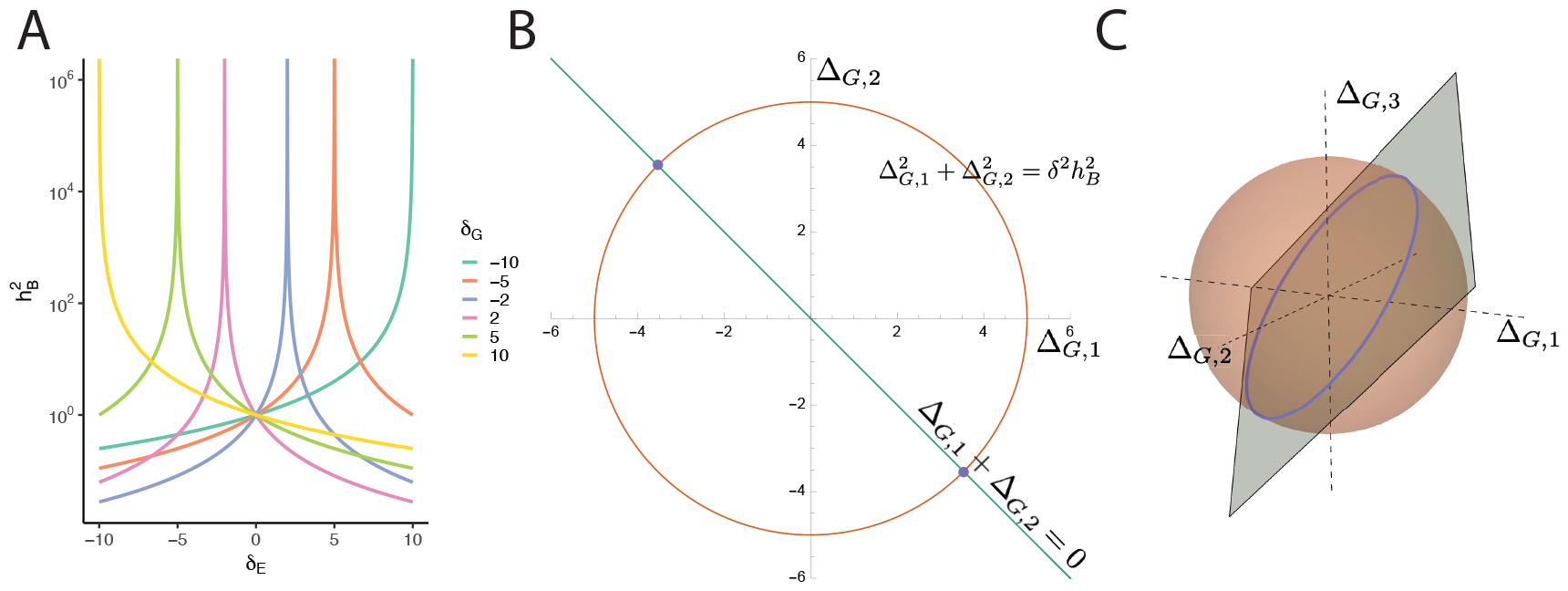
Properties of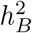. A) The behavior of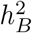 in two populations for different values of the genetic deviation *δ*_*G*_ and environmental deviation *δ*_*E*_. Each curve corresponds to a different genetic deviation. B) Two genetic deviations that are consistent with an observed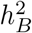with two populations. The circle shows all genetic deviations that hold the genetic variance constant, while the line shows all genetic deviations that sum to 0. The intersection of the line and circle shows the genetic deviations that are consistent with a given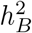. C) An infinite number of genetic deviations consistent with an observed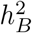 with three populations.The sphere shows all genetic deviations that hold the genetic variance constant, while the plane shows all genetic deviations that result in the genetic deviations summing to 0. The intersection of the sphere and the plane, indicated by the purple circle, contains the values of genetic deviations from the grand mean that will hold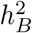 constant.

Finally, the between-group heritability is uninformative about the *direction* of environmental and genetic differences. For the two-population case, this is reflected in the fact that given knowledge of the phenotypic deviation, *δ*, and the between-group heritability, there are two possibile directions of genetic effects that are consistent with the data, namely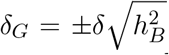. In the Supplementary Text (section S1.3), we show that with more than two groups, there are an infinite number of possible configurations of genetic and environmental deviations that are consistent with a given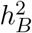. To give some intuition for this result, we start with an alternative representation of the two-group scenario for groups of equal size (Figure 1B). The two axes indicate genetic deviations of the two group means from the grand mean, Δ_*G*,1_ and Δ_*G*,2_. Conditional on a given mean phenotypic difference between the groups, *δ*, obtaining a given value of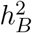 requires that the sum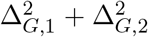 equal a constant, solutions of which are represented by the orange circle. Additionally, because Δ_*G*,1_ and Δ_*G*,2_ are deviations from the grand mean, they must sum to zero, giving the green line corresponding to Δ_*G*,1_ + Δ_*G*,2_ = 0. The two purple points where the line and circle intersect represent the two configurations of the between-group genetic difference that lead to the same value of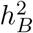.

With a larger number of groups, the situation is entirely analogous, only in higher dimension. Figure 1C shows a schematic of the three-group case. The axes again represent the genetic deviations of each group from the grand mean. The combinations of genetic deviations that results in the same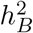 (conditional on phenotypic differences) is now an orange sphere, rather than a circle. And the genetic deviations from the grand mean consistent with the condition that all genetic deviations sum to zero fall on the blue plane (rather than a line). The intersection of the sphere and the plane, shown in purple, gives a circle of genetic deviations on which all have the same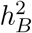 while also being consistent as deviations from a grand mean. With more than three groups, the sphere becomes a hypersphere, the plane becomes a hyperplane, and the same basic relationships hold.

Within-group heritabilities, on their own, provide no information about the causes of between-group differences, in part because any configuration of group means for the genetic contributions to the trait is possible under some value of *r*. The value of *r* itself can be studied under evolutionary models, and in fact it is closely related to the quantity *Q*_*ST*_ studied in evolutionary quantitative genetics, which we consider in the next section.

## *3 P*_*ST*_ analyses can understate or overstate the possibilities for adaptive differentiation

The statistic *Q*_*ST*_ is a measure of the genetic component of trait variation among populations. For diploids, in terms of the variance proportion *r* discussed in the previous section, it is equal to *Q*_*ST*_ = *r/*(2*−r*) (Edge & Rosenberg, 2015). When traits evolve neutrally, *Q*_*ST*_ is expected to be equal to *F*_*ST*_ at putatively neutral sites, which measures the genetic variation among populations (Whitlock, 1999). Thus, by testing whether *Q*_*ST*_ is significantly greater than *F*_*ST*_, it is possible to determine whether there is more genetically explained trait variation among populations than expected under neutral evolution (Leinonen et al., 2013; Prout & Barker, 1993; Spitze, 1993; Whitlock, 2008). Such a pattern is consistent with divergent selection causing trait differences among groups. On the other hand, lower-than-expected values of *Q*_*ST*_ may be taken as evidence of stabilizing selection on a shared optimum.

However, to estimate *Q*_*ST*_ requires arduous common-garden experiments that eliminate environmental confounds. Such common-garden experiments may be difficult or impossible to carry out in some organisms. Thus, it is common in the literature to use a version of *Q*_*ST*_ calculated from phenotypes measured on wild organisms rather than organisms raised in a common garden (for example, Castellani et al., 2023; Salloum et al., 2022; Vitek et al., 2022). This statistic computed from observations of wild organisms, often labeled *P*_*ST*_, is a function of the between-population phenotypic variance, 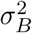, the within-population phenotypic variance, 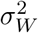, as well as parameters *a* and *b*, which are interpreted as determining the fraction of between- and within-population phenotypic variance that is due to genetics, respectively. For diploids,

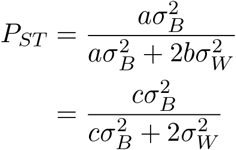

where the parameter *c* = *a/b*. Because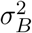and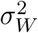 can be estimated from purely phenotypic data but *c* cannot, it is common practice in the literature to explore values of *c* between 0 and 1, seemingly motivated by a suggestion that the genetic component of between-population differences must be proportionally smaller than the genetic component of within-population differences. A well appreciated point is that if the choice of *c* is too large, then *P*_*ST*_ analysis can be anticonservative, leading to spurious inferences of divergent selection. In effect, environmental differences are mistaken for genetic ones, leading to a false rejection of neutral evolution. In the Methods, we show that the coefficients for *P*_*ST*_ are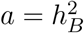, the between-group heritability, and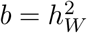, the within-group heritability, so that the correct value of *c* to obtain *P*_*ST*_ = *Q*_*ST*_ is

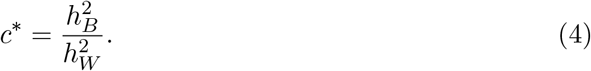

As shown above, 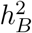 is unconstrained by any measure of genetic and environmental variability within groups. Moreover, because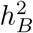 is not constrained to be between 0 and 1, a sensitivity analysis that explores values of *c* between 0 and 1 may lead to extremely conservative tests for divergent selection. Finally, because both *Q*_*ST*_ and *P*_*ST*_ depend on the variance of between-population means, any evolutionary scenarios that produce identical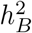will also produce identical *Q*_*ST*_ and *P*_*ST*_, indicating that knowledge of *P*_*ST*_ or *Q*_*ST*_ is not sufficient for disentangling the directions of environmental and genetic effects.

## 4 Simulated example: evolution in the face of an environmental and optimum shift

To demonstrate the impact of our results on evolutionary analyses, we performed populationgenetic simulations using SLiM (Haller & Messer, 2019). In particular, we developed a model in which very different underlying evolutionary forces result in identical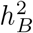 and *Q*_*ST*_, indicating that these statistics are insufficient for characterizing evolutionary history and may be misleading in some circumstances. The scenario we considered is outlined in Figure 2 panels A and D are and inspired by a recent perspective by Harpak and Przeworski (2021): a population evolves with a trait under stabilizing selection until a population split. In one of the populations, the environment and stabilizing selection remain constant; however, in the other population, the environment shifts, along with the optimum trait value. We explore two scenarios: one in which the environment shifts in the same direction as the fitness optimum for the trait (“concordant”, panel A), and one in which they shift in opposite directions (“discordant”, panel D). We use the same optimum shift in both cases, so that the simulations result in a phenotypic difference of *δ* = 1, and set the environmental shift to maintain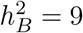at equilibrium.

**Figure 2.**
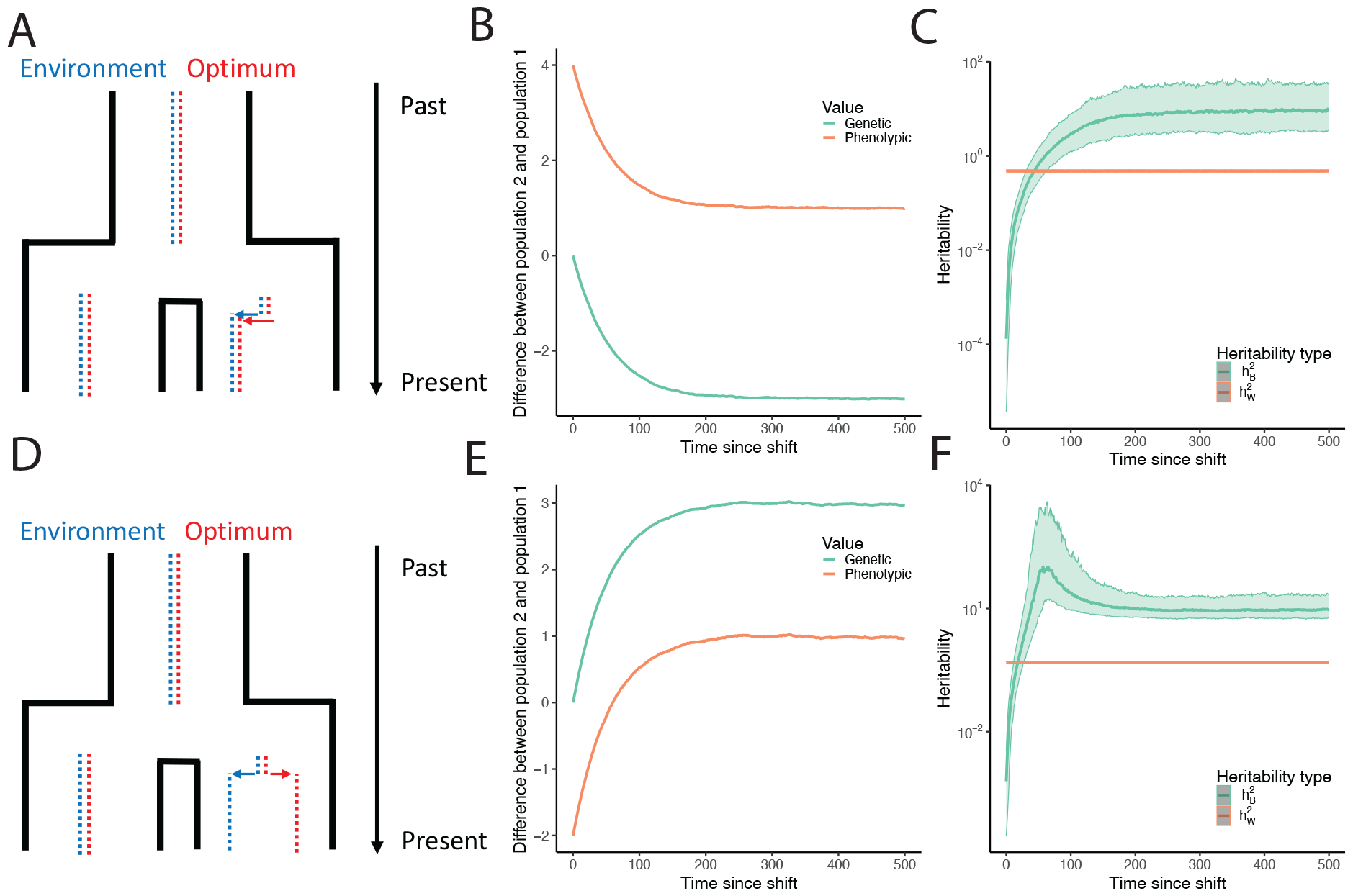
Simulation of a evolutionary scenarios. A, D) Description of the evolutionary scenarios. A population evolves until it splits into two subpopulations of equal size. At the time of split, one of the subpopulations experiences both an environmental shift (in blue) and an optimum shift (red). The environmental shift may either be in the same direction as the optimum shift (concordant, panel A) or in an opposite direction (discordant, panel D). B, E) Evolution of genetic and phenotypic values over time. The horizontal axis shows the number of generations since the split, and the vertical axis shows the value of either the genetic difference between the two populations or the phenotypic difference between the two populations. Concordant shifts are shown in Panel B, discordant shifts in panel E. C, F) Evolution of between- and within-population heritability. The horizontal axis indicates time since the population split, and the vertical axis shows the value of different summaries of phenotypic variation,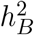and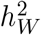; median values are shown with a solid line, and the shaded band indicates the range of values between the 10th and 90th percentiles. Concordant shifts are shown in panel C, discordant shifts in panel F.

Figure 2 panels B and E show the evolution of the difference in phenotypic and genotypic values between the two populations over the course of adaptation. In both directions of environmental shift, the phenotypic differences converge to *δ*_*P*_ = 1, consistent with identical optima shifts in both situations. However, the genetic differences display opposite dynamics, depending on the direction of the environmental shift. If the environmental shift is in the same direction as the optimum shift (panel B), then the adapting population will have a mean genetic value that is less than that of the population that did not experience an environmental shift; in the other scenario (panel E), the adapting population will have a mean genetic contribution to the trait that is greater than the population that did not experience a shift. Thus, to compare the impact of stabilizing selection with environmental change, both the common-garden genetic values *and* the *in situ* phenotypic values are necessary.

Figure 2 panels C and F show that, 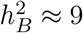at equilibrium by construction. However, the two scenarios approach that value in different ways. When the environment and optimum shift in the same direction, the average approach to equilibrium is monotonic and smooth (panel C). However, when the optimum and environment shift in opposite directions, 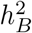 becomes large when the phenotypic difference between the populations is near 0, and comes back down from infinity to settle at the equilibrium value (panel F). It is also much noisier across simulation replicates, as seen by the middle 80% of the distribution spanning 3 orders of magnitude. This is mostly due to the denominator being near zero for a large portion of the simulations (Figure S1). On the other hand, the long-term expected value of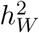 is constant across the simulations and identical regardless of the direction of the environmental shift, as expected, because within-population variability will not change when there is an optimum shift. As emphasized in the previous section, knowledge of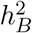at any time during adaptation is not sufficient to characterize the direction in which selective forces are acting.

Finally, we examined *P*_*ST*_ calculations and their impact on the sensitivity of a test to reject neutral trait evolution in these scenarios. In each simulation, we computed *P*_*ST*_ across an array of values of *c* and also computed the true *Q*_*ST*_. Figures 3A and C show the behavior of the statistics and the test of neutral evolution, respectively, when the the environment and optimum trait value shift in the same direction. Here, the behavior of *P*_*ST*_ for a given value of *c* is monotonic, but at different points during the course of adaptation, the same value of *c* can produce either a conservative or anti-conservative estimate of *Q*_*ST*_ from *P*_*ST*_. Early in adaptation, there should be very little genetic signal of adaptation, and *Q*_*ST*_ rejects the null hypothesis approximately 5% of the time at the 5% significance threshold, showing that the test is well calibrated. However, because the environmental shift results in large phenotypic differences, every *P*_*ST*_ test initially results in high false positive detection of adaptation. Subsequently, as adaptation proceeds, *Q*_*ST*_ detects adaptation with high power, although it loses power after approximately 300 generations, because the neutral divergence between populations increases substantially. However, for values of *c <* 1, as typically explored in the literature, *P*_*ST*_ quickly becomes smaller than *Q*_*ST*_, and tests using *P*_*ST*_ fail to reject the null hypothesis significantly earlier into adaptation. Figures 3B and D shows that when the environment and optimum shift in opposite directions, the behavior of *Q*_*ST*_ remains monotonic, but the behavior of *P*_*ST*_ for a given *c* becomes non-monotonic, reaching a minimum when the phenotypic difference between populations is small and then increasing again. This results in complicated behavior of the *P*_*ST*_ test in comparison with the *Q*_*ST*_ test, in which *P*_*ST*_ can go from anti-conservative to conservative, and then back to anti-conservative. Finally, the trajectories of *Q*_*ST*_ are identical regardless of the direction of the environmental shift, in line with the fact that knowledge of *Q*_*ST*_ is not informative of the direction of environmental and genetic shifts.

**Figure 3.**
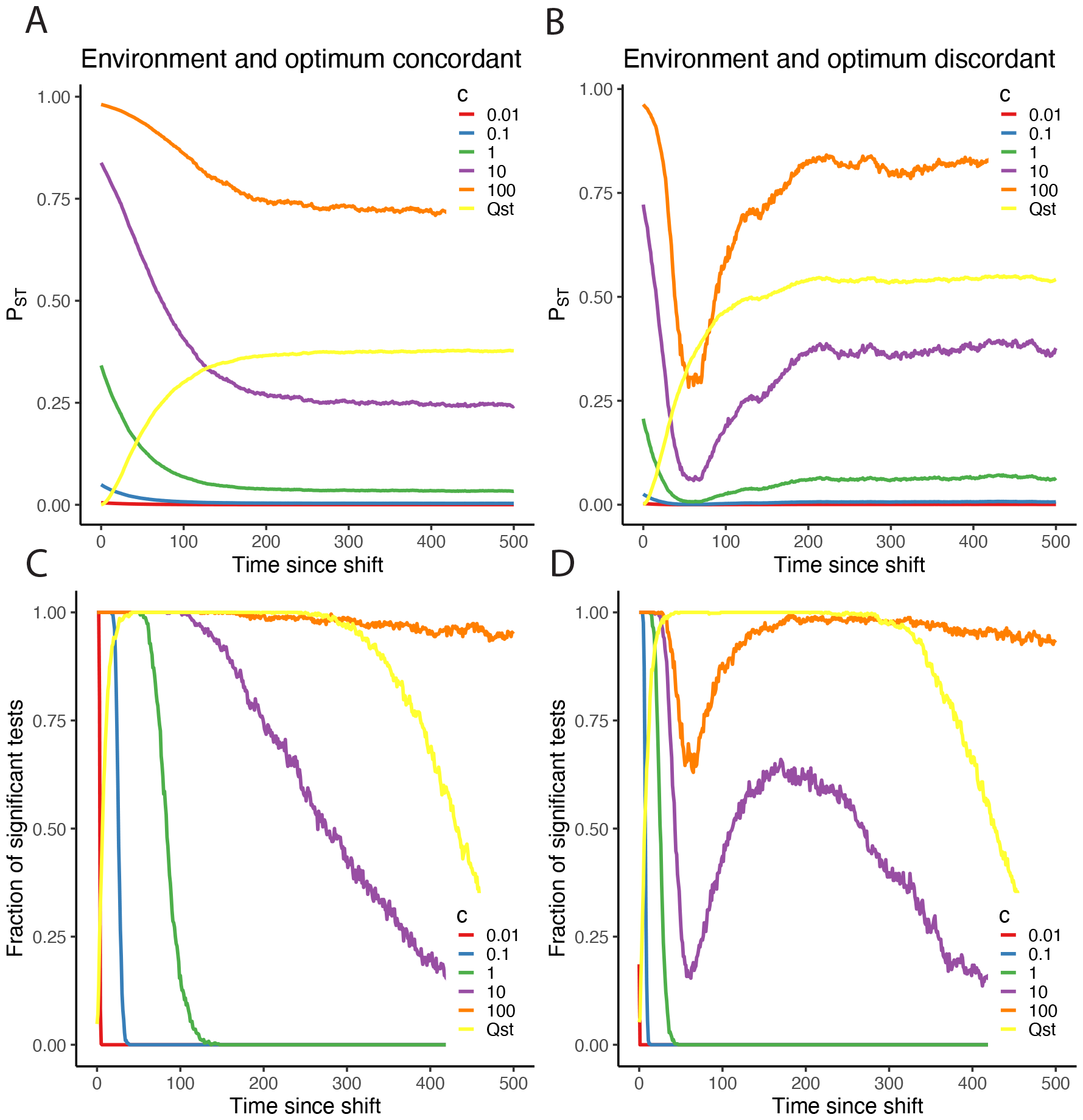
Evolution of *P*_*ST*_ and *Q*_*ST*_. In the top row, the horizontal axis shows the time since the population split, and the vertical axis shows the value of either *Q*_*ST*_ or *P*_*ST*_, with the different colors indicating different values for the choice of *c* in the *P*_*ST*_ formula. In the bottom row, the horizontal axis shows the time since the population split, and the vertical axis shows the fraction of tests in which the null hypothesis of neutral evolution is rejected at the 5% level because *Q*_*ST*_ *> F*_*ST*_. The colors are the same as the top row. A, C) The environment and optimum shifts are concordant. B, D) The environment and optimum shifts are discordant.

## 5 Trait correlations with global ancestry or genome-wide local-ancestry sharing do not reveal the causes of between-group difference

### 5.1 Global ancestry

We have seen that efforts toward quantifying the genetic contribution to differentiation among groups can be confounded by environmental variation. Some have suggested that admixed individuals present opportunities to resolve this difficulty and estimate the genetic component of between-group differences. One suggested approach is to examine the correlation between trait values and global ancestry proportions among admixed individuals (Chakraborty & Weiss, 1986; Chan et al., 2015; Fernández et al., 2003; Kosoy et al., 2012; Molokhia et al., 2003; Rushton & Jensen, 2005). If trait value is positively correlated with the fraction of global ancestry from source population A, then that correlation might be taken as evidence suggesting population A is genetically predisposed to larger trait values than other source populations contributing to the admixed population.

However, this approach depends strongly on assumptions about the relationship between ancestry and environmental effects. We show in the supplementary text (section S.2.1.3) that the expected phenotype of an individual with ancestry fraction *θ*_*i*_ from source population 2 in a two-source admixture is

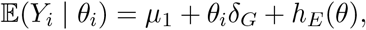

where *μ*_*i*_ is the mean genetic contribution to the trait in population *i* and *h*_*E*_(*θ*) is a function that expresses how the conditional expectation of the environmental effect depends on an individual’s global ancestry fraction. If the environmental and genetic effects of global ancestry are confounded in the admixed population (i.e., if the correlation of *Y* and *h*_*E*_(*θ*) is nonzero), then it may be impossible to recover *δ*_*G*_. As a simple example, consider a case in which individuals with *θ* = 0 experience environmental effects with expectation *E*_1_, individuals with *θ* = 1 experience environmental effects with expectation *E*_2_, and there is a linear gradient between those extremes for individuals of intermediate admixture fraction. If we again define the difference between mean environmental effects to be, *δ*_*E*_ = *E*_2_ *− E*_1_, then *h*_*E*_(*θ*) = *E*_1_ + *θδ*_*E*_, and

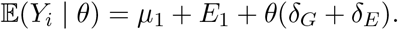

Under this formulation, it would be possible to use linear regression to estimate the slope (in terms of ancestry proportion *θ*) and intercept given a sample of admixed individuals with measured admixture fractions and phenotypes, but the resulting slope would provide information only about the sum of the genetic and environmental effects of global ancestry, not their relative contribution or even their direction.

A similar approach might be applied to admixed siblings, where the correlation with trait value is computed only with respect to variation in realized global ancestry between full siblings, who have the same expected global ancestry proportions on the autosomes. Limiting to within-sibship comparisons would remove some confounds of global ancestry proportion, but it would not exclude environmentally mediated phenotypic differences caused by variation in global ancestry. For example, in African Americans, colorism (Dixon & Telles, 2017) might lead to systematically different treatment as a function of ancestry fraction, even within sets of full siblings.

### 5.2 Local-ancestry heritability

Although variation in global ancestry may be confounded with environmental variation, one might imagine that variation in the genome-wide sharing of local ancestry segments could provide a way forward in identifying the genetic vs. environmental basis of among-group difference.

Zaitlen et al. (2014) proposed a novel estimator of heritability applicable to individuals from admixed populations. The estimator works on principles similar to SNP heritability estimators (Yang et al., 2017), such as implemented in GCTA (Yang et al., 2011). SNP heritability methods work by measuring the degree to which aggregate sharing of alleles at genotyped SNPs among people who are not closely related is correlated with similarity on a phenotype. A strength of SNP heritability methods is that they avoid potential biases stemming from environmental influences shared among closely related people. A weakness is that they cannot inform about contributions to heritability from variants that are not well tagged by SNPs included in the genetic relatedness matrix, in particular rare variants. In local-ancestry heritability estimates, rather than examining shared SNP geno-types, one examines the sharing of segments of local ancestry in an admixed population. For example, consider pairs of individuals from an admixed population formed from two source populations, where both members of all pairs have 50% of their ancestry from each source population. If a trait is heritable, then a pair whose genetic segments from source population 1 fall in exactly the same locations might be expected to be slightly more phenotypically similar than another pair who share no local ancestry segments. Heritability estimates based on local ancestry offer the potential to address one of the main issues facing SNP heritability estimates, since segments of local ancestry tag both common and rare variants. The resulting estimate has also been taken as a proxy for *c* in *P*_*ST*_ analysis (Zaidi et al., 2017), though Zaitlen and colleagues (2014) made no claims about revealing the causes of phenotypic differences among source populations.

Despite the promise of the approach, heritability estimates based on local ancestry face their own challenges. Specifically, directional or stabilizing selection in the history of the source populations can lead to upward or downward biases in heritability estimates, respectively (Huang et al., 2023). In the supplementary text (section S2), we consider this problem and describe a related one that, in a pathological worst case, can lead to heritability estimates near 0 when the true heritability is 1. This problem is most pronounced when the trait under consideration has experienced selection in the source populations, a scenario that Zaitlen et al. (2014) recognized as potentially problematic for their estimator.

In the Supplementary Text (section S2), we decompose the genetic variation for a phenotype in a population having experienced several generations of admixture between two source populations into a component due to variation in global ancestry, a component due to the placement of local ancestry segments conditional on global ancestry, and a component due to randomness in genotype conditional on local ancestry. The variance component principally used by local-ancestry heritability estimation is the variance due to placement of local ancestry segments, conditional on global ancestry,

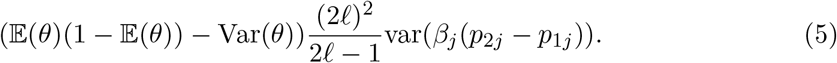

Here, *θ* is a random variable representing global ancestry fraction from source population 2, *ℓ* is the number of (unlinked) loci considered, *β*_*j*_ is the effect size of an effect allele at locus *j, p*_1*j*_ and *p*_2*j*_ are frequencies of the effect allele in populations 1 and 2, and var() indicates the population variance.

Under this model, the difference in genetic means between the source populations is

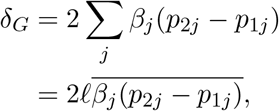

where the overbar denotes the population mean. However, the expression in (5) does not provide a means to estimate *δ*_*G*_, since the average of the quantity *β*_*j*_ (*p*_2*j*_ *− p*_1*j*_) does not appear in equation (5); only its variance does. The mean value of *β*_*j*_ (*p*_2*j*_ *− p*_1*j*_) is not constrained by the variance of the *β*_*j*_ (*p*_2*j*_ *− p*_1*j*_) values. The approach of Zaitlen and colleagues (2014) can be applied with or without a fixed effect for global ancestry fraction. However, comparing these results does not reveal the genetic component of the between-group difference if there may also be environmental effects associated with ancestry fraction, for the reasons discussed in the previous subsection (and see section S2.3).

## 6 Discussion

We showed that knowledge of the heritability of within-population differences does not constrain the heritability of between-population differences at all, as discussed by Lewontin (1970). Our results establish Lewontin’s interpretation generally.

Moreover, even if the between-group heritability were estimable, it is consistent with potentially infinitely many configurations of genetic differences among populations and cannot be used to infer the direction(s) of group differences in genetic contributions to a trait. This second point is especially important. Evolutionary models may be able to identify plausible ranges, under given assumptions, for the variance proportion of the genetic component of the trait accounted for by group membership, *r*, since *r* is a function of *Q*_*ST*_, which is studied in evolutionary quantitative genetics. However, even if *r* (and thus between-group heritability, which has *r* as a key ingredient) can be constrained some-what, the causes of group difference remain ambiguous without a clear understanding of environmental differences among groups and their relevance for the trait. This point was also anticipated by a verbal argument, this one by philosopher Ned Block (1995).

Although our analysis of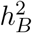was partially anticipated by Lewontin, Block, and others (Kempthorne, 1978; R. C. Lewontin, 1974, 1970; Mackenzie, 1980; Turkheimer, 1991), the existence of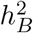 has been invoked for decades, up to the present, to suggest that within-group heritability is informative about among-group differences (Jensen, 1972a, 1972b, 1975; Plomin & DeFries, 1976; Warne, 2020, 2021). As one example, Warne (2021) writes “The existence of the equation [relating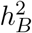to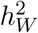] alone shows that the claim that ‘the genetic basis of the difference between two populations bears no logical or empirical relation to the heritability within populations’ (Lewontin, 1970, p. 7) is incorrect.” Our work clarifies that Lewontin’s claim is correct in general and not only in the special cases he used for illustration. The relationship between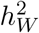 and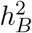 is rather like the relationship between a random variable’s expectation and its variance. One can write the variance as a function of the expectation (i.e. Var(*X*) = E(*X*^2^)*−*E(*X*)^2^), but unless further assumptions are made, knowledge of the expectation alone gives no information about the variance. Roseman & Bird (2023) recently arrived at a similar position, concluding that “there is no relationship between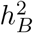 and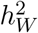 under an additive genetic model…unless evolutionarily explicit models are used to relate them.”

These findings have important implications not just for quantifying the genetic component of among-group differences, but also for methods of detecting genetic adaptation using phenotypic data from multiple populations. As we showed, the phenotypic analogue of *Q*_*ST*_, called *P*_*ST*_, is not constrained by within-population heritability. In fact, because the between-population heritability is not bounded from above by 1, common sensitivity analyses, which assume that the between-population component of genetic differentiation is smaller than the within-population component, can be very conservative.*P*_*ST*_ has been criticized (Brommer, 2011; Pujol et al., 2008) and is often treated cautiously as a result. However, to our knowledge, the fact that *c* can be much larger than 1 is not widely understood, nor is the fact that it can achieve very large values in realistic evolutionary scenarios.

The same fundamental problem that plagues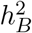and *P*_*ST*_ also prevents the separation of genetic and environmental components of differences among source populations via study of ancestry variation in admixed populations. Variation among admixed people in global ancestry proportions may be associated with both genetic and environmental effects, leading to confounding. Variation among admixed people in the sharing of local ancestry segments provides information that is sometimes useful in heritability estimation, but the resulting estimates do not provide a way forward for estimating differences in the genetic contribution to the trait among source populations, since local-ancestry heritability estimates depend principally on a variance component that is not closely related to the genetic contribution to differences among groups.

All of the phenomena we have discussed so far arise under the simplest possible models, in which phenotypes arise via additive influence of genotype and environment. In the presence of gene-by-environment interaction, the role of genetic factors in explaining group differences itself depends on the environment in which the groups exist. Even careful control of the environment then provides only a partial picture, since it tells us only about the genetic contribution to group difference in the environment studied, which may well change or reverse in a different one (R. C. Lewontin, 1974). For example, Chakraborty & Weiss (1986) studied diabetes incidence in admixed populations with varying proportions of European or Native American ancestries. The increased incidence of diabetes with more Native American ancestries that they observed ran against what seemed to them the most obvious environmental risk factor, the degree to which a group’s diet was westernized. However, as they noted, “earlier prevalence studies did not reveal a high occurrence of [diabetes] in pre-World War II populations of Amerindians” suggesting the inadequacy of a purely genetic model, and a potential role for gene-environment interaction in explaining the observed group differences.

Clearly, our results do not prevent the identification of specific genetic or environmental factors that might contribute to group differences, in the sense that, if their influence were removed, the magnitude or direction of a group difference might change. For example, in domestic dogs and related canid species, a specific variant in *IGF-1* appears to be an important factor in explaining body-size differences (Plassais et al., 2022). This variant is associated with body size within and across species and breeds, in a broad set of different genetic backgrounds and different environments, and it plausibly has a regulatory function that relates it to body size. On the environmental side, exposure to known toxins such as air pollution can lead to health problems such as asthma or some cancers, and exposure is sometimes stratified by race or socioeconomic status, plausibly contributing to health disparities (Evans & Kantrowitz, 2002; Liu et al., 2021). The problems we identify arise when simplistic assumptions are made about aggregate contributions of genetic or environmental differences to group differences, particularly when our understanding of the sources of variation in the phenotype are not understood. In some cases, a single causal factor might be identified, such as a genotype that causes a Mendelian disease, although even such seemingly straightforward cases may be more complicated and susceptible to environmental modification, as in the case of phenylketonuria (Scriver, 2007). But when the causes of variation in a phenotype are poorly understood and most phenotypic variation is unexplained, projection on the basis of known factors is particularly difficult. For example, in humans, attempts to explain group differences in terms of polygenic scores—predicted trait values calculated from genotypes, which are typically noisy predictions—are fraught with difficulties (Coop, 2019; Rosenberg et al., 2018).

When it is possible, the best solution to the problems we point out is careful control of the environment. Common-garden observations support the original uses of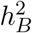 by Lush (1949) and applications of *Q*_*ST*_. Departures from the common garden are unavoidable for studies of humans and of some wild organisms. At the same time, if the environment is not controlled, then the sources of among-group differences become ambiguous, and this ambiguity cannot be resolved by study of within-group variation in aggregate.

If control of the environment is not possible, it may sometimes be possible to observe that a phenotypic difference is persistent across a broad set of environments that vary in all dimensions that might be relevant for the trait. With respect to the methods discussed here, they may allow for firmer conclusions if the environmental sources of trait variation are understood and measured or modeled, even if they cannot be controlled. In practice, it is often difficult to make such conclusions—the relevant environmental variables are unknown, and it is difficult to tell whether they have been explored. In fact, Lewontin (1970) contained a third thought experiment to this effect. In the same case as described in the introduction, with genetic variation within groups but no genetic variation between groups raised in different soils, Lewontin asks us to to imagine a chemist making a survey of the soils. The chemist finds a large difference in nitrate concentrations, but correcting the difference only partially lessens the phenotypic difference between the groups, leading researchers to conclude incorrectly that the remaining phenotypic difference is genetic. In fact, a much subtler difference in zinc levels is “the real culprit.” This thought experiment was inspired by the long delay in the discovery of the importance of trace minerals for plant development, “because ordinary laboratory glassware will leach out enough of many trace elements to let plants grow normally.” Lewontin asked rhetorically, “Should educational psychologists study plant physiology?” We might ask today how we can come to understand the traits we care about well enough to distinguish nitrate cases from zinc cases.

## 7 Methods

### 7.1 A linear model of quantitative traits within and between groups

We model the phenotype of an individual *j* in population *i* as the sum of genetic and environmental components, *G* and *ϵ*, respectively,

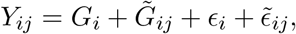

where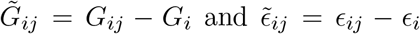 are the genetic and environmental deviations, respectively, of individual *j* from the mean genetic and environmental values for group *i*.

This model is applicable to a trait regardless of genetic architecture, as long as there are no gene-by-environment interactions.

### 7.2 Computing the between- and within-group heritability

In terms of the linear model formulation, the definition o f the b etween-group heritability is

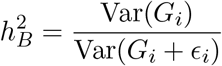

and the within-group heritability is

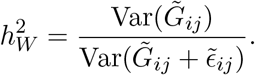

In the Supplementary Text (section S1.2), we show that with these definitions, one can recover the relationship between 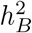 and 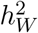 derived by DeFries (1972).

### 7.3 The parameters used in *P*_*ST*_

In terms of the linear model formulation, for diploids, the definition of *Q*_*ST*_ is

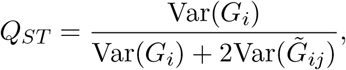

and the definition of *P*_*ST*_ is

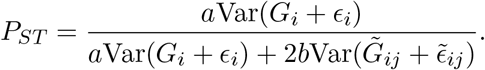

In terms of the variance proportions *t* and *r* in DeFries’ expression for between-group heritability, it follows that *Q*_*ST*_ = *r/*(2 *− r*).

For *P*_*ST*_ = *Q*_*ST*_ we need

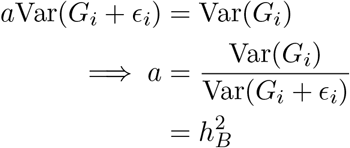

and similarly

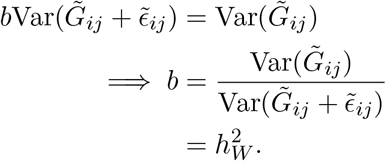

### 7.4 Stabilizing selection simulations

We simulated phenotypes under selection using SLiM (Haller & Messer, 2019). In all simulations, an initial population of 500 diploid individuals with genomes of 1,000,000 positions, with a mutation rate of 10^*−*7^ per generation and a recombination rate of 10^*−*5^ (to approximate free recombination). New mutations have an effect on the trait drawn from a Normal(*μ* = 0,*σ*^2^ = 0.005) distribution. Environmental effects are Gaussian with variance 2. We impose Gaussian stabilizing selection with an optimum at 0 and variance of 100. After 20,000 generations, the populations split into two populations of 20,000 diploid individuals. We then consider two scenarios. In both scenarios, one subpopulation maintains an optimum at 0, while the other experiences an optimum shift to an optimum at 1. In one scenario, the environment shifts to a mean of -2, in the opposite direction of the optimum shift. In the other scenario, the environment shifts to +4, in the same direction as the optimum shift. The simulations are run for 500 generations after the population split. See Table 7.4 for a summary of simulation parameters. For each scenario, we conducted 500 replicate simulations.

**Table 1:** Simulation parameters.

| Scenario | Concordant | Discordant |
| --- | --- | --- |
| N | 500 | 500 |
| Mutation rate | $10^{-7}$ | $10^{-7}$ |
| Recombination rate | $10^{-5}$ | $10^{-5}$ |
| Mutation distribution | N(0,.005) | N(0,.005) |
| Baseline environment distribution | N(0,2) | N(0,2) |
| Baseline selection kernel | N(0,100) | N(0,100) |
| Time of population split | 20,000 | 20,000 |
| Optimum shift | +1 | +1 |
| Environment shift | +4 | -2 |

### 7.5 Testing for adaptation using the Lewontin-Krakauer distribution

To test for adaptation in our simulations by comparing *P*_*ST*_ or *Q*_*ST*_ to *F*_*ST*_, we first computed ratio-of-averages genome-wide *F*_*ST*_ (as implemented in `SLiM’`s `calc_Fst` function) on the basis of neutral markers that were not associated with the trait in the simulations.

The Lewontin-Krakauer (R. C. Lewontin & Krakauer, 1973) test uses the statistic

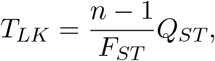

which has *χ*^2^ distribution with *n −* 1 degrees of freedom, where *n* is the number of demes (in our case, *n* = 2). For the tests using *P*_*ST*_, we substituted *P*_*ST*_ for *Q*_*ST*_ in the definition of the Lewontin-Krakauer test statistic. We note that we computed the variance in group means without using Bessel’s correction, as shown by Weaver (2016) to be the correct formula for Nei’s *F*_*ST*_ (Nei, 1973).

## 8 Code availability

Code used in this paper can be found at https://github.com/Schraiber/heritability.

## Acknowledgments

We thank K. Bird, G. Coop, A. Gusev, A. Harpak, M. Pennell, and M. Przeworski for helpful comments on an earlier draft of the manuscript, and members of the Edge, Mooney, and Pennell labs for helpful conversations. We are also grateful for the helpful comments of the anonymous reviewers, which improved the manuscript. We acknowledge support from NIH grant R35GM137758 to MDE.

## Notes

https://github.com/Schraiber/heritability

### Competing Interest Statement

The authors have declared no competing interest.

### Summary of Updates

The figures are updated and improved.

