## Supplementary text for "Heritability within groups is uninformative about differences among groups: cases from behavioral, evolutionary, and statistical genetics"

Compiled on February 11, 2024

In this supplement, we provide further mathematical details supporting the conclusions in the main text. In the below, we adopt a standard quantitative-genetic framework without interactions, in which individual phenotypes are formed by summing a genetic contribution to the phenotype (which we also call a "genetic component" and is also called a "breeding value," "genetic value," or "polygenic effect" in other contexts) and an environmental contribution to the phenotype. In section S1, we discuss between-group heritability as defined in quantitative genetics and adopted in behavior genetics, showing that it underdetermines the genetic differences among groups and is not bounded from above. In section S2, we decompose the phenotypic variation in an admixed population, allowing for some conclusions about methods to estimate heritability using local ancestry variation.

#### S1 Between-group heritability

DeFries (1972), following Lush (1949), expressed the "between-group" heritability as

$$h_B^2 = h_W^2 \frac{(1-t)r}{(1-r)t}, \tag{S1}$$

where $t$ is the intraclass correlation for the phenotype—modeled as a sum of distinct genetic and environmental influences—and $r$ is the intraclass correlation for genetic contributions to the trait in a quantitative genetics framework. Below (section S1.1) we show that even in the simplest case, any given value of $h_B^2$ is attainable under two different arrangements of the underlying parameters, with opposing interpretations. In section S1.2, we build a more general framework, and in section S1.3, we show that this general framework gives rise to a rotational symmetry, such that with more than two groups, there are infinite possible arrangements of the group-level environmental and genetic differences on the phenotype consistent with any given value of the between-group heritability.

#### S1.1 A case of two groups

First, consider the simplest possible case, with two equally sized groups in which the trait is affected by genetic and environmental variation in the same way. For any individual, the effects of genetic and environmental variation are independent and additive, with variances $\sigma_G^2$ and $\sigma_E^2$, so that within-group heritability is $h_W^2 = \sigma_G^2/(\sigma_G^2 + \sigma_E^2)$ in both groups. (This assumption is relaxed in the next subsection.) There are is also a between-group mean difference, $\delta$, with a genetic and environmental component and no interactions, such that $\delta = \delta_G + \delta_E$.

In this setting, the intraclass correlation coefficients are the proportions of variation in the genetic contribution to the trait ($r$) and trait values ($t$) attributable to the group difference. For the genetic components (represented with $G$), this is $r = 1 - (\mathrm{Var}(G|M))/(\mathrm{Var}(G))$, where $M$ denotes group membership, $\mathrm{Var}(G|M) = \sigma_G^2$, and $\mathrm{Var}(G) = \mathbb{E}_M(\mathrm{Var}(G|M)) + \mathrm{Var}_M(\mathbb{E}(G|M)) = \sigma_G^2 + \delta_G^2/4$. (The second term in the last equation is the variance of a Bernoulli(1/2) random variable multiplied by $\delta_G$.) So

$$r = 1 - \frac{\sigma_G^2}{\sigma_G^2 + \delta_G^2/4} = \frac{\delta_G^2}{4\sigma_G^2 + \delta_G^2}.$$

Similarly,

$$t = 1 - \frac{\mathrm{Var}(Y|M)}{\mathrm{Var}(Y)} = \frac{(\delta_G + \delta_E)^2}{4((\sigma_G + \sigma_E)^2) + (\delta_G + \delta_E)^2},$$

where $Y$ is the trait value.

Plugging these values for $r$ and $t$ into Defries' expression for the heritability of the group means (eq. S1) gives

$$h_B^2 = \frac{\sigma_G^2}{\sigma_G^2 + \sigma_E^2} \frac{\left(1 - \frac{(\delta_G+\delta_E)^2}{4(\sigma_G^2+\sigma_E^2)+(\delta_G+\delta_E)^2}\right) \frac{\delta_G^2}{4\sigma_G^2+\delta_G^2}}{\left(1 - \frac{\delta_G^2}{4\sigma_G^2+\delta_G^2}\right) \frac{(\delta_G+\delta_E)^2}{4(\sigma_G^2+\sigma_E^2)+(\delta_G+\delta_E)^2}}$$

Re-expressing the terms in large parentheses,

$$h_B^2 = \frac{\sigma_G^2}{\sigma_G^2 + \sigma_E^2} \frac{\frac{4(\sigma_G^2+\sigma_E^2)}{4(\sigma_G^2+\sigma_E^2)+(\delta_G+\delta_E)^2} \frac{\delta_G^2}{4\sigma_G^2+\delta_G^2}}{\frac{4\sigma_G^2}{4\sigma_G^2+\delta_G^2} \frac{(\delta_G+\delta_E)^2}{4(\sigma_G^2+\sigma_E^2)+(\delta_G+\delta_E)^2}}.$$

The denominators on the right cancel, giving

$$h_B^2 = \frac{\sigma_G^2}{\sigma_G^2 + \sigma_E^2} \frac{4(\sigma_G^2 + \sigma_E^2)\delta_G^2}{4\sigma_G^2(\delta_G + \delta_E)^2}$$

Finally, $4(\sigma_G^2 + \sigma_E^2)\sigma_G^2$ cancels from the numerator and denominator, leaving the simple expression

$$h_B^2 = \frac{\delta_G^2}{(\delta_G + \delta_E)^2} = \frac{\delta_G^2}{\delta^2}$$

In some ways, this expression is intuitive, in that $\delta_G^2$ is proportional to the variance in the group means due to genetic differences, and $(\delta_G + \delta_E)^2 = \delta^2$ is proportional to the total variance in group means. However, the interpretation as a proportion is misleading. Assuming nonzero values of $h_B^2$ and the group difference $\delta$, there are always two values of $\delta_G$ possible, namely $\delta_G = \pm\sqrt{h_B^2\delta^2}$. This means that the heritability of the group mean is the same regardless of whether the genetic group difference points in the same direction as the group difference or in the opposite direction. Concretely, if the observed group difference is 10 in favor of group 1, $h_B^2$ is equal to 0.25 if either the genetic group difference is +5 for group 1 (and the environmental difference is thus also +5 for group 1), or the genetic group difference is -5 for group 1 (and the environmental difference is +15 for group 1).

It is also possible to obtain values of $h_B^2 > 1$ (and, in fact, arbitrarily large $h_B^2$ values) if the genetic and environmental components of the group difference point in opposite directions. In general, the interpretation of $h_B^2$ only makes sense if the environmental and genetic components of the group differences agree in sign. Even then, it is a somewhat odd statistic in the two-group case: in the case that the differences point in the same direction, the square root of $h_B^2$, that is, $\delta_G/\delta$, would be more direct.

The interpretive difficulties can be thought of as emerging in part from the application of $h_B^2$ to a two-group framework, which is different from Lush's scenario, which considered the groups as many pens of animals. In particular, the typical assumption of heritability methods that environmental and genetic group differences are uncorrelated cannot manifest in actual instantiations of the two group case. The differences will either point in the same direction, or in opposite directions. However, we show below that larger numbers of groups do not solve the ambiguities.

#### S1.2 A linear-model formulation

Next, we reformulate the same model as in the previous subsection as a linear model, allowing easy extension to larger numbers of groups. Consider a set of phenotype values, where $Y_{ij}$ is the phenotype value of the $j$th individual from the $i$th subgroup. We decompose the phenotype values into (additive) genetic effects ($G_{ij}$) and environmental effects ($\epsilon_{ij}$), as is standard.

$$Y_{ij} = G_{ij} + \epsilon_{ij}$$

We then rewrite $G_{ij}$ as the sum of the group-mean genetic component in group $i$ and individual $j$'s deviation from the mean of their group,

$$G_{ij} = G_i + \tilde{G}_{ij},$$

where $G_i$ is the mean genetic component of individuals in group $i$ and $\tilde{G}_{ij} = G_{ij} - G_i$. Similarly, for the environmental component,

$$\epsilon_{ij} = \epsilon_i + \tilde{\epsilon}_{ij}.$$

Then we have

$$Y_{ij} = G_i + \tilde{G}_{ij} + \epsilon_i + \tilde{\epsilon}_{ij}.$$

The typical expression for (narrow-sense) heritability is, in this notation,

$$h^2 = \frac{\mathrm{Var}(G_{ij})}{\mathrm{Var}(Y_{ij})}.$$

The between-group heritability is

$$h_B^2 = \frac{\mathrm{Var}(G_i)}{\mathrm{Var}(G_i + \epsilon_i)}$$

The within-group heritability is

$$h_W^2 = \frac{\mathrm{Var}(\tilde{G}_{ij})}{\mathrm{Var}(\tilde{G}_{ij} + \tilde{\epsilon}_{ij})}, \tag{S2}$$

In general, this is not necessarily equal to any of the within-group heritabilities or to their arithmetic average, though it is equal to both of these if the within-group heritabilities are the same in each group.

The intraclass genetic correlation is

$$r = \frac{\mathrm{Var}(G_i)}{\mathrm{Var}(G_{ij})}$$

and the intraclass phenotypic correlation is

$$t = \frac{\mathrm{Var}(G_i + \epsilon_i)}{\mathrm{Var}(Y_{ij})}.$$

Thus,

$$\begin{aligned} h_B^2 &= \frac{\mathrm{Var}(G_i)}{\mathrm{Var}(G_i + \epsilon_i)} \\ &= \frac{\mathrm{Var}(G_i)}{\mathrm{Var}(G_i + \epsilon_i)} \frac{\mathrm{Var}(G_{ij})}{\mathrm{Var}(G_{ij})} \frac{\mathrm{Var}(Y_{ij})}{\mathrm{Var}(Y_{ij})} \\ &= \frac{\mathrm{Var}(G_i)}{\mathrm{Var}(G_{ij})} \frac{\mathrm{Var}(Y_{ij})}{\mathrm{Var}(G_i + \epsilon_i)} \frac{\mathrm{Var}(G_{ij})}{\mathrm{Var}(Y_{ij})} \\ &= \frac{r}{t} h^2, \end{aligned}$$

where $h^2$ is the "total" heritability—the variance of the genetic component over the variance of the phenotype across all groups. This is the expected result. A similar calculation gives the within-group heritability, as expressed in equation (S2), as

$$h_W^2 = \frac{1-r}{1-t} h^2.$$

Combining these results shows that

$$h_B^2 = h_W^2 \frac{r}{t} \frac{1-t}{1-r},$$

as in DeFries and in equation (S1) above.

To proceed, we need to work out some of the variances, generally using the law of total variance to condition on group membership, $i = m$, where $1 \leq m \leq M$ (i.e. $M$ total groups). We then assume that a fraction $p_m$ samples are from group $m$.

$$\begin{aligned} \mathrm{Var}(G_{ij}) &= \mathbb{E}(\mathrm{Var}(G_{ij}|i=m)) + \mathrm{Var}(\mathbb{E}(G_{ij}|i=m)) \\ &= \mathbb{E}(\sigma_{G,m}^2) + \mathrm{Var}(\mu_{G,m}) \\ &= \sum_{m=1}^{M} p_m \sigma_{G,m}^2 + \sum_{m=1}^{M} p_m (\mu_{G,m} - \bar{\mu}_G)^2, \end{aligned}$$

where $\sigma_{G,m}^2 \equiv \mathrm{Var}(\tilde{G}_{ij}|i=m)$, $\mu_{G,m}$ denotes the group-mean genetic component, and $\bar{\mu}_G$ is the grand-mean genetic component, $\sum_{m=1}^{M} p_m \mu_{G,m} = \bar{\mu}_G$.

A similar calculation gives the variance of the phenotype value as

$$\mathrm{Var}(Y_{ij}) = \sum_{m=1}^{M} p_m \sigma_{Y,m}^2 + \sum_{m=1}^{M} p_m (\mu_{Y,m} - \bar{\mu}_Y)^2,$$

where substituting $Y$ for $G$ indicates analogous descriptions of phenotypic values as opposed to genetic contributions to phenotype (for example, $\sigma_{Y,m}^2 \equiv \mathrm{Var}(\tilde{G}_{ij} + \tilde{\epsilon}_{ij}|i=m)$).

Analysis of this model yields the two-group result in the previous subsection as a special case. Starting from the definition of $h_B^2$ directly,

$$h_B^2 = \frac{\mathrm{Var}(G_i)}{\mathrm{Var}(G_i + \epsilon_i)}.$$

we only need to plug in the appropriate variances. Using the same notation as above,

$$\mathrm{Var}(G_i) = \sum_{m=1}^{M} p_m (\mu_{G,m} - \bar{\mu}_G)^2.$$

In the case of two groups of equal size considered in the previous subsection, $M = 2$, $\mu_{G,2} - \mu_{G,1} = \delta_G$ implies $\mu_{G,i} - \bar{\mu}_G = \delta_G/2$, and $p_m = 1/2$, so

$$\begin{aligned}\sum_{m=1}^{M} p_m(\mu_{G,m} - \bar{\mu}_G)^2 &= \sum_{m=1}^{2} \frac{1}{2}(\delta_G/2)^2 \\ &= \frac{1}{4}\delta_G^2.\end{aligned}$$

A similar calculation on the full phenotypic variance, noting that the case of two groups of equal size implies $\mu_{P,i} - \bar{\mu}_P = (\delta_G + \delta_E)/2$, yields

$$\mathrm{Var}(G_i + \epsilon_i) = \frac{1}{4}(\delta_G + \delta_E)^2.$$

Thus, we arrive at the expected result,

$$h_B^2 = \frac{\delta_G^2}{(\delta_G + \delta_E)^2}.$$

A notable feature of this derivation is that the within-group genetic and phenotypic variances are never used, and no assumptions are made about them. Thus, $h_B^2$ is not constrained by the within-group heritabilities or their components.

#### S1.3 Symmetry for multiple groups

For multiple groups, we obtain a strong symmetry result. To simplify notation, let

$$\Delta_{G,i} = \mu_{G,i} - \bar{\mu}_G$$

i.e. the difference between the mean genetic effect in group $i$ and the overall mean genetic effect. Similarly,

$$\Delta_{Y,i} = \mu_{Y,i} - \bar{\mu}_Y.$$

where we can decompose $\Delta_{Y,i} = \Delta_{G,i} + \Delta_{E,i}$, where $\Delta_{E,i}$ denotes a group-level deviation from the grand mean in the environmental effect on the trait. Note that in contrast to the values $\delta_G$, etc. defined in the previous sections as the difference between two populations, the values $\Delta_G$, etc., used here indicate deviations from the grand mean. In the case of two populations with equal size, $\Delta_{G,1} = \Delta_{G,2} = \frac{1}{2}\delta_G$, for example. To simplify the argument, we assume here that the groups are all of equal size, but it can be generalized to groups of unequal size.

The between-group heritability is

$$
\begin{aligned}
h_B^2 &= \frac{\mathrm{Var}(G_i)}{\mathrm{Var}(Y_i)} \\
&= \frac{\sum_{m=1}^{M}(\mu_{G,m} - \bar{\mu}_G)^2}{\sum_{m=1}^{M}(\mu_{Y,m} - \bar{\mu}_Y)^2} \\
&= \frac{\sum_{m=1}^{M}\Delta_{G,m}^2}{\sum_{m=1}^{M}\Delta_{Y,m}^2} \\
&= \frac{\|\Delta_G\|^2}{\|\Delta_Y\|^2},
\end{aligned}
$$

where the last line follows by writing the differences into the elements of two vectors, $\Delta_G = (\Delta_{G,1}, \ldots \Delta_{G,M})^T$ and $\Delta_P = (\Delta_{P,1}, \ldots \Delta_{P,M})^T$ and using $\|\cdot\|$ to denote the magnitude of a vector.

Now, suppose we have a matrix $A \in O(M)$, the M-dimensional orthogonal group. The orthogonal group preserves distances—that is, if $\|\mathbf{x}\| = r$ for some M-dimensional vector $\mathbf{x}$, then $\|A\mathbf{x}\| = r$ for $A \in O(M)$. Concretely, this means that $A^T A = I_M$. However, we need a subgroup of $O(M)$, because the fact that the $\delta$ terms are deviations from the grand mean imposes the additional constraint that

$$
\begin{aligned}
\Delta_{G,1} + \Delta_{G,2} + \ldots + \Delta_{G,M} &= 0 \\
\implies \Delta_G^T e &= 0
\end{aligned}
$$

where $e = \frac{1}{\sqrt{M}}(1, 1, \ldots, 1)^T$ is the vector of $M$ 1's, scaled to magnitude 1.

To obtain a transformation that satisfies both requirements, we first pick a matrix to rotate the coordinate system to align a dimension (say, the $M$th) with the vector $e$; call the matrix that does this transformation $P^T \in O(M)$. This maps $(1, 1, \ldots, 1)^T$ to $(0, 0, \ldots, 1)^T$ and $(\delta_1, \delta_2, \ldots, \delta_n)^T$ to $(\delta_1', \delta_2', \ldots, 0)^T$. Now we wish to rotate such that we preserve the length of the vector but also preserve the final axis. This can be done with an axis-preserving rotation,

$$
A = \begin{bmatrix} Q & \mathbf{0}_{M-1} \\ \mathbf{0}_{M-1}^T & 1 \end{bmatrix},
$$

where $\mathbf{0}_n$ is the vector of $n$ 0's and $Q \in O(M-1)$. Note that $A \in O(M)$, because

$$\begin{aligned}
A^T A &= \begin{bmatrix} Q & \mathbf{0}_{M-1} \\ \mathbf{0}_{M-1}^T & 1 \end{bmatrix}^T \begin{bmatrix} Q & \mathbf{0}_{M-1} \\ \mathbf{0}_{M-1}^T & 1 \end{bmatrix} \\
&= \begin{bmatrix} Q^T & \mathbf{0}_{M-1} \\ \mathbf{0}_{M-1}^T & 1 \end{bmatrix} \begin{bmatrix} Q & \mathbf{0}_{M-1} \\ \mathbf{0}_{M-1}^T & 1 \end{bmatrix} \\
&= \begin{bmatrix} Q^T Q & \mathbf{0}_{M-1} \\ \mathbf{0}_{M-1}^T & 1 \end{bmatrix} \\
&= I_M
\end{aligned}$$

Then, we build the transformation by $\Delta_G \to PAP^T \Delta_G$. Then we have that $PAP^T \in O(M)$ (since all $P, A \in O(M)$), but also

$$\begin{aligned}
(PAP^T \Delta_G)^T e &= (\Delta_G^T P A^T P^T) e \\
&= (\Delta_G^T P) A^T (P^T e) \\
&= \Delta_G^T P P^T e \\
&= \Delta_G^T e.
\end{aligned}$$

The third line follows because $P^T e$ is an eigenvector of $A$ with eigenvalue 1, by construction of $A$.

This shows that $h_B^2$ is invariant under a group of transformations that is isomorphic to $O(M-1)$. Thus, for $M = 2$, there are 2 possible vectors of group-mean genetic components, and for all $M > 2$, there are infinitely many possible group-mean genetic-value vectors that are consistent with a given phenotype and genotype.

Note that to hold $\|\Delta_Y\|^2$ constant implies that the environmental differences are rotated the same amount as the genetic differences, which can be seen by picking two different rotations, say $R$ and $S$, to rotate the genetic and environmental components, respectively, obtaining a new phenotype vector, $\Delta_Y'$. Then

$$\begin{aligned}
\|\Delta_Y'\|^2 &= \|R\Delta_G + S\Delta_E\|^2 \\
&= \|R\Delta_G\|^2 + \|S\Delta_E\|^2 + (R\Delta_G)^T (S\Delta_E) \\
&= \|\Delta_G\|^2 + \|\Delta_E\|^2 + \Delta_G^T R^T S \Delta_E.
\end{aligned}$$

The third line follows by construction of the matrices $R$ and $S$. To have this equal the magnitude of the original $\Delta_P$ vector, we need $R^T S = I$. By uniqueness of the inverse, this shows that $R = S$, so one must rotate both components by the same amount.

### S2 Admixed populations

#### S2.1 Genetic variance in an admixed population

We derive the genetic variance (i.e., the variance of the genetic contribution to a trait) in an admixed population formed from two source populations. We use a model of $\ell$ unlinked autosomal loci and assume that a diploid individual with global ancestry fraction $\theta$ has exactly $2\ell\theta$ alleles from source population 2, and $2\ell(1-\theta)$ alleles from population 1. We further assume that conditional on $\theta$, which alleles are from population 1 vs. 2 is decided as a random permutation. Thus, the number of alleles from population 2 at a given locus is approximately Binomial(2,$\theta$) if $\ell$ is not small, and there is no correlation in ancestry across sites conditional on global ancestry. These assumptions do not hold in the case of very recent admixture—for example, individuals with a parent from each source assignment have exactly one allele of each ancestry at every locus—but they should hold approximately after several generations of random mating in an admixed population. They also hold if we consider two distinct populations with no admixture, so that each individual has either $\theta = 0$ or $\theta = 1$.

A similar derivation appears in a recent preprint from Huang et al. (2023), who first derive the variance in genotype at a single locus and then sum across all loci. We take a distinct approach because we are interested in separating the variance into three components: a component due to randomness in genotype conditional on local ancestry, a component due to the specific configuration of local ancestry conditional on global ancestry, and a component due to variation in global ancestry. Although we arrive at similar expressions, our results differ because we assume that the global ancestry exactly equals the sum of the local ancestry of the loci, whereas Huang et al. (2023) draw local ancestry from a binomial distribution parameterized by the global ancestry. The difference between our expressions is small for large numbers of loci, being proportional to a factor of $\ell/(\ell-1)$, where $\ell$ is the number of loci, as we show below.

Define $\gamma_{ij} \in \{0, 1, 2\}$ the number alleles with ancestry from source population 2 at locus $j$ in individual $i$. Define $p_{1j}$ and $p_{2j}$ the frequency of an "effect" allele in ancestries 1 and 2 at locus $j$. Define $\beta_j$ the effect on a trait of the effect allele at locus $j$, which we assume to be the same regardless of global or local ancestry of an individualz. The difference between the source populations in mean genetic contribution to the trait is

$$\delta_G = 2\sum \beta_j(p_{2j} - p_{1j}). \tag{S3}$$

$\theta_i$ is the global ancestry of individual $i$, and $\theta_i = \sum_{j=1}^{\ell} \gamma_{ij}/(2\ell)$.

We want the variance of the genetic contribution to the phenotype, which we label $Y_i$, or $\mathrm{Var}(Y_i) = \mathrm{Var}(\sum \beta_j G_{ij})$, where $G_{ij}$ is the genotype of individual $i$ at locus $j$. Using the law of total variance,

$$\mathrm{Var}(Y_i) = \mathbb{E}_{\theta_i}(\mathrm{Var}(Y_i|\theta_i)) + \mathrm{Var}_{\theta_i}(\mathbb{E}(Y_i|\theta_i)). \tag{S4}$$

We will break the first term in equation S4 into two pieces by using the law of total variance once again on the conditional variance, giving

$$\mathrm{Var}(Y_i) = \mathbb{E}_{\theta_i}(\mathbb{E}_{\gamma|\theta_i}(\mathrm{Var}(Y_i|\gamma_{i1}, ..., \gamma_{i\ell}; \theta_i) + \mathrm{Var}_{\gamma|\theta_i}(\mathbb{E}(Y_i|\gamma_{i1}, ..., \gamma_{i\ell}; \theta_i))) + \mathrm{Var}_{\theta_i}(\mathbb{E}(Y|\theta_i)). \tag{S5}$$

Now we have three pieces,

- $\mathbb{E}_\theta(\mathbb{E}_{\gamma|\theta}(\mathrm{Var}(Y_i|\gamma_{i1}, ..., \gamma_{i\ell}; \theta_i)))$, the variance due to random genotypes conditional on local ancestry
- $\mathbb{E}_\theta(\mathrm{Var}_{\gamma|\theta}(\mathbb{E}(Y_i|\gamma_{i1}, ..., \gamma_{i\ell}; \theta_i))$, the variance due to random placement of local ancestry segments conditional on global ancestry
- $\mathrm{Var}_{\theta_i}(\mathbb{E}(Y|\theta_i))$, the variance due to randomness in global ancestry proportion.

##### S2.1.1 Genetic variance due to randomness in genotype conditional on local ancestry

We start with the first term in equation S5, the variation due to randomness in genotype conditional on local ancestry. Starting with the term inside the expectation,

$$\mathrm{Var}(Y_i|\gamma_{i1}, ..., \gamma_{i\ell}; \theta_i) = \mathrm{Var}\left(\sum \beta_j G_{ij}|\gamma_{i1}, ..., \gamma_{i\ell}; \theta_i\right).$$

Genotypes at distinct sites are uncorrelated conditional on local ancestry, so we can split into three sums corresponding to the three possible local ancestry values at each locus,

$$\begin{aligned}\mathrm{Var}(Y_i|\gamma_{i1}, ..., \gamma_{i\ell}; \theta_i) = &\sum_{j:\gamma_{ij}=0} 2\beta_j^2 p_{1j}(1-p_{1j}) \\ &+ \sum_{j:\gamma_{ij}=1} \beta_j^2[p_{1j}(1-p_{1j}) + p_{2j}(1-p_{2j})] \\ &+ \sum_{j:\gamma_{ij}=2} 2\beta_j^2 p_{2j}(1-p_{2j}).\end{aligned}$$

That is, every allele drawn from population 1 at locus $j$ contributes $\beta_j^2 p_{1j}(1-p_{1j})$ to the conditional variance, and every allele drawn from population 2 contributes $\beta_j^2 p_{2j}(1-p_{2j})$. Because proportion $\theta_i$ of the alleles have ancestry from source population 2, we can write

.

$$\mathbb{E}_{\gamma|\theta_i}(\mathrm{Var}(Y_i|\gamma_{i1}, ..., \gamma_{i\ell}; \theta_i)) = 2\sum_j \beta_j^2[(1-\theta_i)p_{1j}(1-p_{1j}) + \theta_i p_{2j}(1-p_{2j})]. \tag{S6}$$

Finally, we take the expectation of the right side of eq. S6 over variation in global ancestry $\theta$, leading to an expression that matches eq 1.1 of Huang et al. (2023),

$$\mathbb{E}_\theta(\mathbb{E}_{\gamma|\theta}(\mathrm{Var}(Y_i|\gamma_{i1}, ..., \gamma_{i\ell}; \theta_i)) = 2\sum_j \beta_j^2[(1-\mathbb{E}(\theta))p_{1j}(1-p_{1j}) + \mathbb{E}(\theta)p_{2j}(1-p_{2j})]. \quad \text{(S7)}$$

###### S2.1.2 Genetic variance due to random placement of local ancestry conditional on global ancestry

Next, we analyze the second component of the genetic variance as expressed in equation S5, that due to the placement of local ancestry segments conditional on global ancestry, or $\mathbb{E}_{\theta_i}(\mathrm{Var}_{\gamma|\theta_i}(\mathbb{E}(Y_i|\gamma_{i1}, ..., \gamma_{i\ell}; \theta_i))$. Starting from the middle,

$$\mathbb{E}(Y_i|\gamma_{i1}, ..., \gamma_{i\ell}; \theta_i) = \mathbb{E}\left(\sum_j \beta_j G_{ij} \mid \gamma_{i1}, ..., \gamma_{i\ell}; \theta_i\right),$$

where $G_{ij} \in \{0, 1, 2\}$ is the number of effect alleles at locus $j$. Because the probability of an effect allele depends on the local ancestry, we can write this as a sum of three terms,

$$\mathbb{E}(Y_i|\gamma_{i1}, ..., \gamma_{i\ell}; \theta_i) = \sum_{j:\gamma_{ij}=0} 2\beta_j p_{1j} + \sum_{j:\gamma_{ij}=1} \beta_j[p_{1j} + p_{2j}] + \sum_{j:\gamma_{ij}=2} 2\beta_j p_{2j}.$$

To make the next step explicit, we will define some new notation to convert the diploid version of this problem into a haploid one—that is, treating the diploid sets of $\ell$ loci as a set of $2\ell$ haploid alleles—simplifying the final expressions. Define $\zeta_{ik} \in \{0, 1\}$ with $k \in \{1, ..., 2\ell\}$ as an alternative representation of the local ancestry of individual $i$. Specifically, if $\zeta_{ik} = 1$ for $k \leq \ell$, then at locus $k$, diploid individual $i$ has ancestry from population 2 on their first haplotype, and $\zeta_{ik} = 0$ indicates ancestry from population 1 in the same position. For $k > \ell$, $\zeta_{ik} = 1$ indicates that individual $i$ has ancestry from population 2 at locus $k - \ell$ on their second haplotype. Thus, $\zeta$ simply represents local ancestry as a set of $2\ell$ indicator variables, and random placement of local ancestry corresponds to a random permutation of the $\zeta$ values. Similarly, define $\alpha_1, ..., \alpha_{2\ell}$ to be equal to the concatenation of two copies of the effect sizes $\beta_1, ..., \beta_\ell$, and define $q_{m1}, ..., q_{m(2\ell)}$ to be the concatenation of two copies of $p_{m1}, ..., p_{m\ell}$ for $m \in \{1, 2\}$. Then

$$\mathbb{E}(Y_i|\zeta_{i1}, ..., \zeta_{i(2\ell)}; \theta_i) = \sum_{k=1}^{2\ell} (\alpha_k q_{1k} + \zeta_{ik}\alpha_k(q_{2k} - q_{1k})).$$

We need the variance of this term with respect to variation in local ancestry placement. The first term does not depend on the local ancestry encoding, so

$$\mathrm{Var}_{\gamma|\theta_i}(\mathbb{E}(Y_i|\zeta_{i1}, ..., \zeta_{i(2\ell)}; \theta_i)) = \mathrm{Var}_{\gamma|\theta_i}\left(\sum_{k=1}^{2\ell} \zeta_{ik}\alpha_k(q_{2k} - q_{1k})\right). \quad \text{(S8)}$$

Because variation in $\gamma$ given $\theta$ corresponds to random permutation of the $\zeta$ values under our assumptions, this is simply the variance of the sum of $2\ell\theta_i$ values of $\alpha_k(q_{2k} - q_{1k})$ drawn from $2\ell$ possibilities without replacement. The standard formula for the variance of a sum of $n$ values obtained by simple random sampling from a population of $N$ objects with population variance $\sigma^2$ is

$$\frac{N-n}{N-1} n\sigma^2,$$

where the term $(N-n)/(N-1)$ is sometimes called a finite population correction (Bondy and Zlot 1976). Substituting $2\ell\theta_i = n$ and $2\ell = N$ gives

$$\mathrm{Var}_{\gamma|\theta}(\mathbb{E}(Y_i|\zeta_{i1}, ..., \zeta_{i(2\ell)}; \theta_i)) = \theta_i(1-\theta_i)\frac{(2\ell)^2}{2\ell-1}\sigma^2, \tag{S9}$$

where $\sigma^2$ is the (population) variance of the products $\alpha_k(q_{2k} - q_{1k})$, which, by construction, is equal to the variance of the per-locus products of effect size and allele-frequency difference, $\beta_j(p_{2j} - p_{1j})$. Finally, we need the expectation with respect to global ancestry of this conditional variance,

$$\begin{aligned}\mathbb{E}_\theta(\mathrm{Var}_{\gamma|\theta}(\mathbb{E}(Y_i|\gamma_{i1}, ..., \gamma_{i\ell}; \theta_i))) &= \mathbb{E}_\theta[\theta_i(1-\theta_i)\frac{(2\ell)^2}{2\ell-1}\mathrm{var}(\beta_j(p_{2j}-p_{1j}))] &\text{(S10)}\\ &= (\mathbb{E}(\theta)(1-\mathbb{E}(\theta)) - \mathrm{Var}(\theta))\frac{(2\ell)^2}{2\ell-1}\mathrm{var}(\beta_j(p_{2j}-p_{1j})), &\text{(S11)}\end{aligned}$$

where var() denotes the population variance. This can also be written as

$$(\mathbb{E}(\theta) - \mathbb{E}(\theta^2))\frac{(2\ell)^2}{2\ell-1}\mathrm{var}(\beta_j(p_{2j}-p_{1j})).$$

###### S2.1.3 Genetic variance due to variation in global ancestry

Finally, we analyze the third component of the genetic variance as expressed in equation S5, that due variation in global ancestry, or $\mathrm{Var}_\theta(\mathbb{E}(Y|\theta))$.

We first compute the conditional expectation of the genetic component given the global ancestry fraction. We can express this using the law of total expectation in terms of local ancestry at each locus,

$$\begin{aligned}\mathbb{E}(Y_i \mid \theta_i) &= \mathbb{E}_{\gamma|\theta_i}(\mathbb{E}(Y_i \mid \gamma_{i1}, \ldots, \gamma_{il}; \theta_i) \mid \theta_i)\\ &= \mathbb{E}_{\gamma|\theta_i}\left(\mathbb{E}\left(\sum_j \beta_j G_j \mid \gamma_{i1}, \ldots, \gamma_{il}; \theta_i\right) \mid \theta_i\right).\end{aligned}$$

Separating loci according to local ancestry, we have

$$\mathbb{E}(Y_i \mid \gamma_{i1}, \ldots, \gamma_{il}; \theta_i) = \sum_{j:\gamma_{ij}=0} 2\beta_j p_{1j} + \sum_{j:\gamma_{ij}=1} \beta_j \left(p_1 + p_2\right) + \sum_{j:\gamma_{ij}=2} 2\beta_j p_{2j},$$

because an allele drawn from population 1 at locus $j$ contributes $\beta_j p_{1j}$ to the mean, and similarly for alleles drawn from population 2. Finally, because an allele comes from population 1 with probability $1 - \theta_i$ and from population 2 with probability $\theta_i$, we have

$$\mathbb{E}(Y_i \mid \theta_i) = \sum_j 2\beta_j \left(p_{1j} + \theta(p_{2j} - p_{1j})\right) = \mu_1 + \theta_i \delta_G, \tag{S12}$$

where $\delta_G = \mu_2 - \mu_1 = \sum_j 2\beta_j(p_{2j} - p_{1j})$ is the difference in genetic means between groups. Thus,

$$\mathrm{Var}_\theta(\mathbb{E}(Y|\theta)) = \mathrm{Var}_\theta(\mu_1 + 2\theta\delta_G.) = \delta_G^2 \mathrm{Var}(\theta). \tag{S13}$$

We can then write

$$\begin{aligned}
\delta_G^2 &= \left(\sum 2\beta_j(p_{2j} - p_{1j})\right)^2 \\
&= 4\left(\sum(\beta_j(p_{2j} - p_{1j}))^2 + \sum_{j \neq k} \beta_j\beta_k(p_{2j} - p_{1j})(p_{2k} - p_{1k})\right),
\end{aligned}$$

so that

$$\mathrm{Var}_\theta \mathbb{E}(Y|\theta)) = 4\mathrm{Var}(\theta)\left(\sum(\beta_j(p_{2j} - p_{1j}))^2 + \sum_{j \neq k} \beta_j\beta_k(p_{2j} - p_{1j})(p_{2k} - p_{1k})\right). \tag{S14}$$

.

It follows from equation (S12) and linearity of expectation that if there are environmental effects associated with ancestry, then the conditional expectation of the trait becomes

$$\mathbb{E}(Y_i \mid \theta_i) = \sum_j 2\beta_j \left(p_{1j} + \theta(p_{2j} - p_{1j})\right) = \mu_1 + \theta_i\delta_G + h_E(\theta_i),$$

where $h_E(\theta)$ is a function giving the conditional expectation of the environmental effect given the ancestry proportion $\theta$. In the special case in which $h_E(\theta) = m + \theta\delta_G$, with $m$ a constant, the variance in $Y$ explained by the linear effect of global ancestry fraction becomes

$$\mathrm{Var}_\theta(\mathbb{E}(Y|\theta)) = (\delta_G + \delta_E)^2 \mathrm{Var}(\theta). \tag{S15}$$

###### S2.1.4 Combining the components

Thus, the variance of the genetic contribution to the trait in an admixed population is obtained as in equation S5, plugging in the values of the three terms as evaluated in equations S7, S10, and S13,

$$\begin{aligned}
\mathrm{Var}(Y) = 2\sum_j \beta_j^2[(1-\mathbb{E}(\theta))p_{1j}(1-p_{1j}) + \mathbb{E}(\theta)p_{2j}(1-p_{2j})] \quad &(S16)\\
+ (\mathbb{E}(\theta)(1-\mathbb{E}(\theta)) - \mathrm{Var}(\theta))\frac{(2\ell)^2}{2\ell-1}\mathrm{var}(\beta_j(p_{2j}-p_{1j})) \quad &(S17)\\
+ \delta_G^2\mathrm{Var}(\theta). \quad &(S18)
\end{aligned}$$

Or, alternatively, expressing the group difference in mean genetic contribution $\delta_G$ in terms of the underlying effect sizes and allele-frequency differences as in equation S14,

$$\begin{aligned}
\mathrm{Var}(Y) = 2\sum_j \beta_j^2[(1-\mathbb{E}(\theta))p_{1j}(1-p_{1j}) + \mathbb{E}(\theta)p_{2j}(1-p_{2j})] \quad &(S19)\\
+ (\mathbb{E}(\theta)(1-\mathbb{E}(\theta)) - \mathrm{Var}(\theta))\frac{(2\ell)^2}{2\ell-1}\mathrm{Var}(\beta_j(p_{2j}-p_{1j})) \quad &(S20)\\
+ 4\mathrm{Var}(\theta)\left(\sum_j(\beta_j(p_{2j}-p_{1j}))^2 + \sum_{j\neq k}\beta_j\beta_k(p_{2j}-p_{1j})(p_{2k}-p_{1k})\right). \quad &(S21)
\end{aligned}$$

This expression is similar to the one in Huang and colleagues (2023) eq. 1.1-1.4. One superficial difference is that our terms are grouped differently, so that they correspond to the variance due to randomness in genotypes (equation S19), local ancestry placement (equation S20), and global ancestry (equation S21). Another superficial difference comes from notation: we use $\theta$ to denote proportion of ancestry from source population 2, whereas they use it for the proportion from source population 1.

However, a more substantive difference arises due to our term

$$\frac{(2\ell)^2}{2\ell-1}\mathrm{Var}(\beta_j(p_{2j}-p_{1j}))$$

from equation S10. To make our formula equivalent to that of Huang et al. (2023), one must make the approximation

$$\frac{(2\ell)^2}{2\ell-1}\mathrm{Var}(\beta_j(p_{2j}-p_{1j})) \approx \sum_j(\beta_j^2(p_{2j}-p_{1j})^2).$$

The difference arises due to our assumption that the genome is of finite length. Recall that to go from equation (S8) to (S9), we used a formula for the variance of a sum of samples from a finite population, and constrained the sum over local ancestry to be equal to the global ancestry. Instead, we can directly apply the rules for the variance of a sum of random variables (recalling that in this case, the local ancestry $\zeta_{ik}$ is random but the effect size, $\beta_j$, and allele frequencies, $p_{1j}$ and $p_{2j}$, are fixed) to obtain

$$\begin{aligned}\mathrm{Var}_{\gamma|\theta_i}\left(\sum_{k=1}^{2\ell}\zeta_{ik}\alpha_k(q_{2k}-q_{1k})\right) =&\sum_{k=1}^{2\ell}\mathrm{Var}(\zeta_{ik})(\alpha_j(q_{2j}-q_{1j}))^2 \\ &+\sum_{1\leq j<k\leq 2\ell}\mathrm{Cov}(\zeta_{ij},\zeta_{ik})\alpha_j(q_{2j}-p_{1j})\alpha_k(q_{2k}-p_{1k})\end{aligned}$$

The constraint that local ancestries at a finite number of loci sum to a fixed global ancestry, $\theta_i$ induces a negative correlation between the local ancestry at different loci equal to

$$\mathrm{Cov}(\zeta_{ij},\zeta_{ik}) = -\frac{\theta_i(1-\theta_i)}{2\ell-1},$$

which, after plugging into the formula above and manipulating, produces our result. On the other hand, if the genome is infinitely long or if the local ancestries are not constrained to add up to the global ancestry, the covariance terms vanish, and one obtains the result from Huang et al. (2023).

The two formulations will be approximately equal when the number of loci, $\ell$, is large, such that $(2\ell)^2/(2\ell-1)\approx 2\ell$ **and** the mean of $\beta_j(p_{2j}-p_{1j})$ across loci is small, i.e. $\frac{1}{\ell}\beta_j(p_{2j}-p_{1j})\approx 0$. The latter condition obtains when the mean genetic difference between source populations is near zero, but it can also obtain for non-zero mean differences between source populations as long as the number of loci is large enough. However, if only one of these conditions is satisfied, there may still be substantial deviation between the two formulas.

#### S2.2 Basis for heritability estimation

The basis for the heritability estimator from Zaitlen et al. (2014) can be seen by making assumptions and applying approximations to the expressions in the previous subsection. After making these approximations, the genetic variance due to local ancestry (Equation (S20)) divided by the total genetic variance will be equal to $F_{ST}$. Thus, dividing an estimate of Equation (S20) by $F_{ST}$ will yield an estimate of the total genetic variance. Similarly, dividing an estimate of the heritability accounted for by local ancestry by $F_{ST}$ will yield an estimate of the heritability in the admixed population.

To see this, first make the approximation outlined in the previous subsection, so that

$$\frac{(2\ell)^2}{2\ell-1}\mathrm{Var}(\beta_j(p_{2j}-p_{1j}))\approx\sum_j(\beta_j^2(p_{2j}-p_{1j})^2).$$

We also must make the assumption that the difference in mean genetic contribution to the trait between the two ancestral populations is 0, i.e. $\delta_G = 0$. Then,

$$\delta_G^2 = \sum_j (\beta_j (p_{2j} - p_{1j}))^2 + \sum_{j \neq k} \beta_j \beta_k (p_{2j} - p_{1j})(p_{2k} - p_{1k}) = 0.$$

Finally, we must assume that there is *no* variance in global ancestry, i.e.

$$\mathrm{Var}(\theta) = 0$$

Under these assumptions, the genetic variance becomes

$$\mathrm{Var}(Y) = 2 \sum_j \beta_j^2 [(1 - \mathbb{E}(\theta)) p_{1j} (1 - p_{1j}) + \mathbb{E}(\theta) p_{2j} (1 - p_{2j})] \tag{S22}$$

$$+ 2(\mathbb{E}(\theta)(1 - \mathbb{E}(\theta))) \sum_j \beta_j^2 (p_{2j} - p_{1j})^2 \tag{S23}$$

$F_{ST}$ in the admixed population is the variance proportion due to ancestry in an indicator variable representing a single draw of an allele from the population. Using the law of total variance, $F_{ST}$ can be written as

$$F_{ST} = \frac{\mathbb{E}(\theta)(1 - \mathbb{E}(\theta)) \sum (p_{2j} - p_{1j})^2}{\mathbb{E}(\theta)(1 - \mathbb{E}(\theta)) \sum (p_{2j} - p_{1j})^2 + (1 - \mathbb{E}(\theta)) \sum p_{1j}(1 - p_{1j}) + \mathbb{E}(\theta) \sum p_{2j}(1 - p_{2j})}. \tag{S24}$$

Note that $F_{ST}$ would be equal to Equation (S23) divided by the sum of equations (S22) and (S23) if the factor of $\beta_j^2$ were not in each summand in Equations (S22) and (S23). Nonetheless, if $\beta_j$ were independent of the allele-frequency difference between populations, then it could be pulled out of the sum, so that

$$\sum_j \beta_j^2 (p_{2j} - p_{1j})^2 = \left( \sum_j \beta_j^2 \right) \left( \sum_j (p_{2j} - p_{1j})^2 \right).$$

Thus, if an estimator of Equation (S23) is available, multiplying it by the inverse of $F_{ST}$ will cancel out the Equation (S23) term, producing

$$\left( \sum_j \beta_j^2 \right) \Big( \mathbb{E}(\theta)(1 - \mathbb{E}(\theta)) \sum (p_{2j} - p_{1j})^2 + (1 - \mathbb{E}(\theta)) \sum p_{1j}(1 - p_{1j}) + \mathbb{E}(\theta) \sum p_{2j}(1 - p_{2j}) \Big).$$

This expression is equal to the genetic variance expressed in equations (S22-S23) if, in addition to being independent of the allele-frequency differences, the values of $\beta_j^2$ are uncorrelated with the sum of the allelic variances in each population, such that they can be pulled out of equation S22. Dividing the estimated genetic variance by the phenotypic variance produces a heritability estimate.

Zaitlen et al. (2014) divide the estimated variance due to local ancestry by $2\theta(1-\theta)F_{ST}$ rather than $F_{ST}$, but this is because they use a non-standard version of $F_{ST}$. Further, in humans, the average value of $\beta^2$ is known to depend on allele frequency. To address this, Zaitlen and colleagues perform their estimation in allele-frequency bins and take a weighted average of the bin-wise estimates.

The rationale presented here for the Zaitlen et al. (2014) estimator is based on the idea that the variance-components estimator of genetic variance due to local ancestry (typically as implemented in GCTA) has the quantity in equation S23 as its estimand. However, we showed that some assumptions are required to eliminate other contributions to the variance, including that the difference in genetic means between the source populations is zero, and that there is no variance in global ancestry. Additionally, one needs to assume that (at least within allele-frequency classes) squared effect sizes are approximately uncorrelated with allele-frequency differences and with the weighted sum of the allelic variances within the source populations. Whereas in expectation there will be no difference in mean genetic trait values between source populations if genetic drift is the source of population differentiation, this expectation is unlikely to hold exactly (Whitlock 1999; Edge and Rosenberg 2015). Similarly, although variance in global ancestry will asymptotically approach zero after many generations, it is unlikely to be zero in most realistic admixed populations. Nonetheless, the variance in ancestry fraction is empirically small in many situations. For example, in the ASW population of African Americans, Huang and colleagues (2023) estimate that $\mathbb{E}(\theta)(1-\theta) \approx 0.178$, whereas $\mathrm{Var}(\theta) \approx 0.018$. Thus, as confirmed in Huang et al. (2023), the estimator of genetic variance in an admixed population proposed by Zaitlen et al. (2014) works in limited circumstances.

Finally, the impact of adding a fixed effect for global ancestry can be extremely pronounced under certain circumstances. In a pathological worst case, consider a trait for which $\mathrm{Var}(\beta_j(p_{2j}-p_{1j})) = 0$, but $\sum_j \beta_j^2(p_{2j}-p_{1j})^2$ is large (for intuition, imagine that the trait is the global ancestry fraction itself, though the condition is somewhat more general than this.) In such a case, any effect of ancestry is entirely due to global ancestry—the specific placement of ancestry segments does not matter, since the effect of local ancestry on the trait is the same at every locus. Thus, if a fixed effect for global ancestry is included, the estimated heritability is $\approx 0$, even if the true heritability is 1. If the fixed effect is excluded, the estimate of heritability will be substantially larger, but it generally will not be accurate either.

##### S2.3 Implications for genetic variance estimation

The expression for the total genetic variance in an admixed population in equations S19-S21 has some important implications for estimation of genetic variance in an admixed population, as discussed recently by Huang and colleagues (2023). For example, SNP heritability estimates, such as in GCTA (Yang et al. 2011), cannot capture variance due to the second term in equation S21. This second term is large if polygenic selection has driven group mean genetic components of the trait apart, and can be negative if groups are experiencing stabilizing selection with the same optimum (Latta 1998; Berg and Coop 2014), leading to under- or over-estimation of the genetic variance, and thus of the heritability in the admixed population. The fact that the genetic variance due to local ancestry cannot take account of the second term in S21 also means that it cannot serve on its own as a basis for estimating the group mean difference in the genetic contribution to the trait, $\delta_G$.

In practice, Zaitlen and colleagues (2014) include a fixed effect for individual global ancestry. This fixed effect might in principle be used to get information about $\delta_G$, and it is in fact straightforward to use it to estimate $\delta_G$ if it can be assumed that the environmental contribution to the trait is independent of global ancestry. But if the environment is not controlled, it is also possible that there is a difference in mean environmental contribution to the trait in the two source populations (in our notation, $\delta_E \neq 0$). That difference may also translate to the admixed population, and one possibility is that it does so in a linear way, so that an individual's expected environmental component is $\mu_{E,1} + \theta_i \delta_E$. In this case, $\delta_G$ in equation S13 is replaced by $\delta_G + \delta_E = \delta$, the mean phenotypic difference between the source populations. Thus, the genetic difference is confounded with the environmental difference and cannot be identified separately using the fixed effect of global ancestry. There may be strategies for separating the environmental and genetic components if the form of the relationship between environmental effects and genetic ancestry is known with some precision and differs from the genetic effects, or if environmental effects of ancestry are constant within sibships, allowing analysis of differences in realized genetic ancestry among siblings. But in general, it may be difficult or impossible to say whether such conditions hold without the ability to control the environment. Further, if there are both genetic and environmental effects of global ancestry, then there is gene-environment correlation in the admixed population, and the definition of heritability in the admixed population becomes ambiguous. Similar conceptual questions arise if genetic effects of global ancestry cause systematic environmental variation that then influences the phenotype, analogously to Jencks' famous thought experiment about red hair in a society that denies schooling to red-headed children (Jencks 1972).

Our focus in this paper is on possibilities for separating $\delta_G$ and $\delta_E$, genetic and environmental contributions to group mean difference under a fully additive model. Local-ancestry heritability is not a viable approach—the random effect associated with local ancestry is, under some assumptions, connected to the variation due to random genotypes conditional on local ancestry, but it is not closely connected to $\delta_G$. The genetic contribution to the group difference $\delta_G$ appears in the fixed effect of global ancestry, but it can be perfectly confounded with $\delta_E$, so there is no general basis for estimating $\delta_E$ separately. Thus, although local-ancestry heritability has been used as a proxy for the parameter $c$ in $P_{ST}$ analysis (Zaidi et al. 2017), it cannot specify the correct value of $c$. Whether the $c$ estimates that result have useful properties in tests of selection is a matter for separate investigation.

#### S3 Supplementary Figure

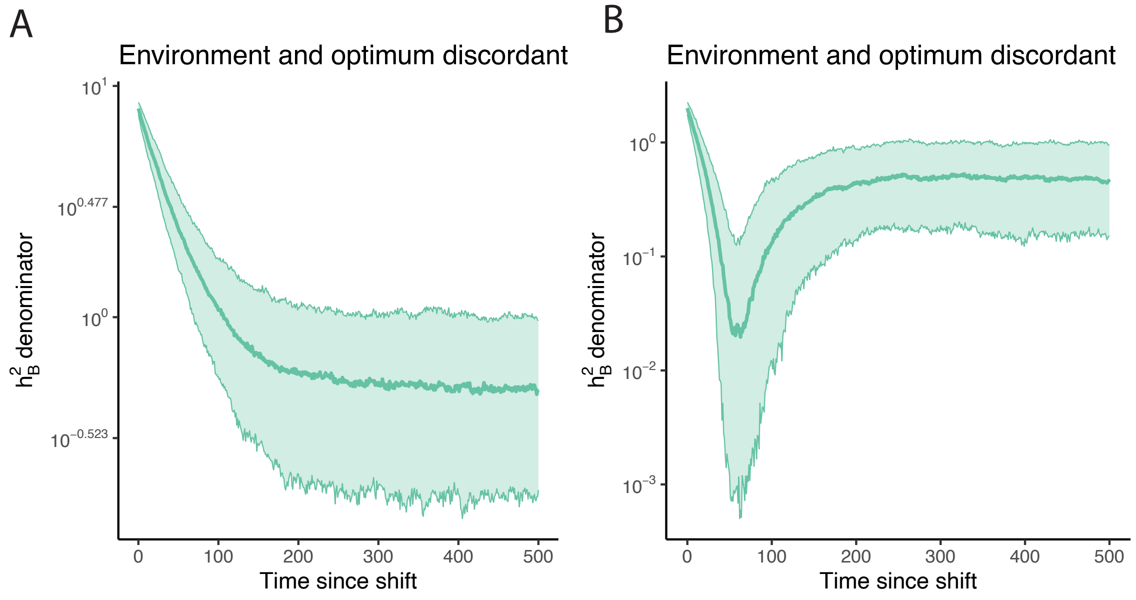


Figure S1: The denominator of $h_B^2$. In each panel, the horizontal axis shows the time since the population split and the vertical axis shows the value of the denominator of $h_B^2$. The median is shown with a solid line and values between the 10th and 90th percentiles are shaded. A) The environment and optimum shifts are concordant. B) The environment and optimum shifts are discordant.
